# Structure and Activity of the Essential UCH Family Deubiquitinase DUB16 from *Leishmania donovani*

**DOI:** 10.1101/2025.02.28.640783

**Authors:** James A. Brannigan, Mohd Kamran, Nathaniel G. Jones, Elisa M. Nightingale, Eleanor J. Dodson, Sarfaraz A. Ejazi, Jeremy C. Mottram, Nahid Ali, Anthony J. Wilkinson

**Affiliations:** York Structural Biology Laboratory, York Biomedical Research Institute and Department of Chemistry, University of York, York YO10 5DD, UK; Infectious Diseases and Immunology Division, CSIR-Indian Institute of Chemical Biology, Kolkata, West Bengal 700032, India; York Biomedical Research Institute and Department of Biology, University of York, York YO10 5DD, UK

**Keywords:** deubiquitinase, UCH family, substrate specificity, Leishmania, crystal structure

## Abstract

In Leishmania parasites, as for their hosts, the ubiquitin proteasome system is important for cell viability. As part of a systematic gene deletion study, it was discovered that four cysteine protease type deubiquitinases (DUBs) are essential for parasite survival in the promastigote stage, including DUB16. Here we have purified and characterised recombinant DUB16 from *Leishmania donovani*, which belongs to the ubiquitin C-terminal hydrolase (UCH) family. DUB16 efficiently hydrolyses C-terminal aminocoumarin and rhodamine conjugates of ubiquitin consistent with proposed cellular roles of UCH-type DUBs in regenerating free monomeric ubiquitin from small molecule ubiquitin adducts arising from adventitious metabolic processes. The crystal structure of DUB16 reveals a typical UCH-type deubiquitinase fold, and a relatively short and disordered crossover loop that appears to restrict access to the catalytic cysteine. At close to stoichiometric enzyme to substrate ratios, DUB16 exhibits deubiquitinase activity towards diubiquitins linked through isopeptide bonds between Lys11, Lys48 or Lys63 residues of the proximal ubiquitin and the C-terminus of the distal ubiquitin. With 100-1000-fold higher turnover rates, DUB16 cleaves the ubiquitin-ribosomal L40 fusion protein to give the mature products. A DUB-targeting cysteine-reactive cyanopyrrolidine compound, IMP-1710, inhibits DUB16 activity. IMP-1710 was shown in promastigote cell viability assays to have parasite killing activity with EC_50_ values of 1−2 μM, comparable to the anti-leishmanial drug, miltefosine. *L. mexicana* parasites engineered to overproduce DUB16 showed a modest increase in resistance to IMP-1710, providing evidence that IMP-1710 inhibits DUB16 *in vivo*. Together these results suggest on-target activity and that DUB16 may be a druggable target to develop new anti-leishmania compounds.

## Introduction

Leishmaniasis is a complex disease caused by more than 20 different leishmania parasite species which are transmitted by sand flies. It ranges in severity from cutaneous (CL) to visceral (VL) forms (1, 2). More than 1 billion people live in areas where leishmaniasis is endemic and it is estimated there are up to 1 million and 30,000 new cases of CL and VL respectively each year (https://www.who.int/health-topics/leishmaniasis). There is no vaccine available for use in humans and the drugs currently used to treat leishmaniasis are toxic, expensive or lose effectiveness over time as drug resistance develops (2, 3). There is thus a pressing need for new drugs and although there are a number of promising pre-clinical candidates, new targets are needed to sustain the drug discovery pipeline (4) (5).

The ubiquitin proteasome system is a promising therapeutic target for the treatment for leishmaniasis (6–8) with the Novartis inhibitor LXE408 undergoing phase II clinical trials. As a result, we have been exploring the wider ubiquitination system in these parasites to identify and provide genetic and chemical validation of new anti-leishmanial drug targets (9–11). Ubiquitin (Ub) is a highly conserved 76 amino acid residue protein with a β-grasp fold whose attachment to proteins alters their fate in complex ways (12). Ubiquitination involves the formation of isopeptide bonds between the C-terminus of Ub and lysine side chains of acceptor proteins. It is brought about by the sequential action of Ub-activating enzymes (E1s), Ub-conjugating enzymes (E2) and Ub-ligases (E3s). Substrate proteins can be monoubiquitinated at one or multiple sites. In many instances, further Ubs are attached to one of the seven lysine side chains giving rise to polyubiquitination. In addition, linear ubiquitin chains can be formed because of conventional peptide bonds between the C-terminal glycine and the N-terminal methionine.

For ubiquitination in *Leishmania mexicana*, it is predicted that two E1 ubiquitin activating enzymes feed 13 E2 ubiquitin conjugating enzymes which serve 79 E3 ubiquitin ligases comprising 61 RING (really interesting new gene), 12 HECT (homologues to E6-AP C-terminus) five U-box and one RBR (RING-between-RING) type enzymes (13). In trypanosomatids, attachment of ubiquitin and ubiquitin-like proteins (Ubls) to target proteins regulates the cell cycle, endocytosis, protein sorting and degradation, autophagy and various aspects of infection and stress responses (9). Ubiquitination is reversed by the action of deubiquitinases (DUBs). DUBs have been classified into multiple families of cysteine proteases and a family of zinc-metalloproteases (14). In *L. mexicana*, there are 20 genes encoding cysteine protease-type deubiquitinases (10). Seventeen belong to the ubiquitin-specific protease (USP) family, two, DUB15 and DUB16, belong to the ubiquitin C-terminal hydrolase (UCH) family and the remaining one, DUB17, belongs to the ovarian tumour protease (OTU) family. Six of these, DUBs 2, 15, 16, 17, 18 and 19, were detected in *L. mexicana* cell lysates by profiling with an activity-based probe (10). In a systematic gene deletion study, null mutants of DUBs 1, 2, 12 and 16 could not be generated in promastigotes in the absence of an episomal copy of these genes, providing evidence that these enzymes are essential (15). For DUB2, inducible deletion by the DiCre system provided additional evidence of essentiality. Meanwhile, 10 of the remaining 16 DUBs analysed are needed variously for parasite development to the amastigote form, infection and survival in the mammalian host (10).

To return ubiquitin to the cellular pool, the DUBs must collectively recognise and cleave linkages of various types (14). Some deubiquitinases exhibit specificity for a particular linkage type (16) while others exhibit little specificity towards their substrate (17). Besides their role in reversing the action of the E3 ligases, deubiquitinases are needed for the post-translational processing of ubiquitin precursors. In the genomes of many species, ubiquitin is encoded as a fusion to the L40 protein from the 60S ribosomal subunit, or as a variable and often large number of tandem ubiquitin repeats – over 40 in many *Leishmania* species (18).

The interest here is in the UCH family deubiquitinase, DUB16 which is present and active in *L. mexicana* promastigotes (10). The human host has four UCH-type deubiquitinases with varying tissue distributions and disease associations. UCHL1 is very highly expressed in neuronal tissues and its dysfunction is associated with neurodegenerative disease (19). Dysfunction of all four UCH-type human DUBs is associated with the progression of a multitude of cancer types prompting detailed investigations of their structure and function and their targeting for drug discovery (20, 21).

Despite their importance, relatively little is known of the substrate preferences of the UCH family members. Early studies showed these enzymes were general hydrolases that regenerate ubiquitin from small amide and thiol derivatives formed adventitiously (22). The size restriction was later attributed to the presence of a crossover loop that appears to limit substrate access to the active site (23–26). UCH family members have however elsewhere been shown to cleave specific Ub-fusion protein substrates including Ub-ribosome precursors and Ub fusions to Ube2W and SUMO2 (27, 28) but they are not able to cleave lysine linked or linear ubiquitin dimers (29, 30).

With a view to understanding its essential role in parasite viability and further validating DUB16 as a drug target for leishmaniasis, we have expressed and purified recombinant DUB16 from *L. donovani* and determined its substrate specificity in deubiquitination assays and its 3D structure by X-ray crystallography.

## Results

### Parasite and host deubiquitinases

DUB16 from *L. donovani* (Gene ID LdBPK_250190.1), is a 233 residue (25 kDa) protein that is predicted to comprise a single catalytic domain belonging to the UCH family. There are four UCH-type DUBs in the human host UCHL1, UCHL3, UCHL5 (also known as UCH37) and BAP1 (BRCA1-associated protein) with their DUB domains exhibiting pairwise sequence identities of 24−55 % (**Supplementary Table S1**). The *L. donovani* enzymes, DUB15 and DUB16, have 26 % identity. Unlike DUB16, DUB15 is dispensable in *L. mexicana* promastigotes (10). Their closest human homologues are UCHL3 (37 %) and UCHL5 (42 %) respectively (**Supplementary Table S1**). For DUB15 and UCHL5, the sequence identity extends into the C-terminal helical domain, which for the human orthologue, binds to the proteasome subunit RPN13 and the NFRKB subunit of the INO80 complex which mediates ATP-dependent nucleosome sliding (31).

### Purification and characterisation of DUB16

The coding sequence of DUB16 was amplified from *L. donovani* genomic DNA and ligated into a vector directing the overproduction of polyhistidine-tagged protein in *E. coli* (**Supplementary Table 2**). The enzyme was purified in high yield (∼50 mg per litre of bacterial cell culture) by tandem steps of immobilised metal affinity chromatography, with an intervening treatment with human rhinovirus 3C protease (HRV 3C), followed by size exclusion chromatography. DUB16 was judged to have over 95% purity by Coomassie staining of SDS polyacrylamide gels.

### Activity against ubiquitin-small molecule conjugates

The activity of purified DUB16 was first assayed with a fluorogenic substrate, Z-RLRGG-AMC, in which the C-terminal five residues of ubiquitin (Arg-Leu-Arg-Gly-Gly), have a carboxybenzyl modified N-terminus and 7-amino-4-methylcoumarin linked at the C-terminus. Cleavage of the C-terminal amide bond leads to a strong enhancement of the coumarin fluorescence. The time courses of the fluorescence increase at substrate concentrations in the range 0-100 μM allowed a k_cat_ of 2 +/− 0.2 s^−1^ and a K_M_ of 36 +/− 7 μM to be determined from a fit to the Michaelis-Menten equation (**Figure 1A; Supplementary Figure S1A**).

**Figure 1.**
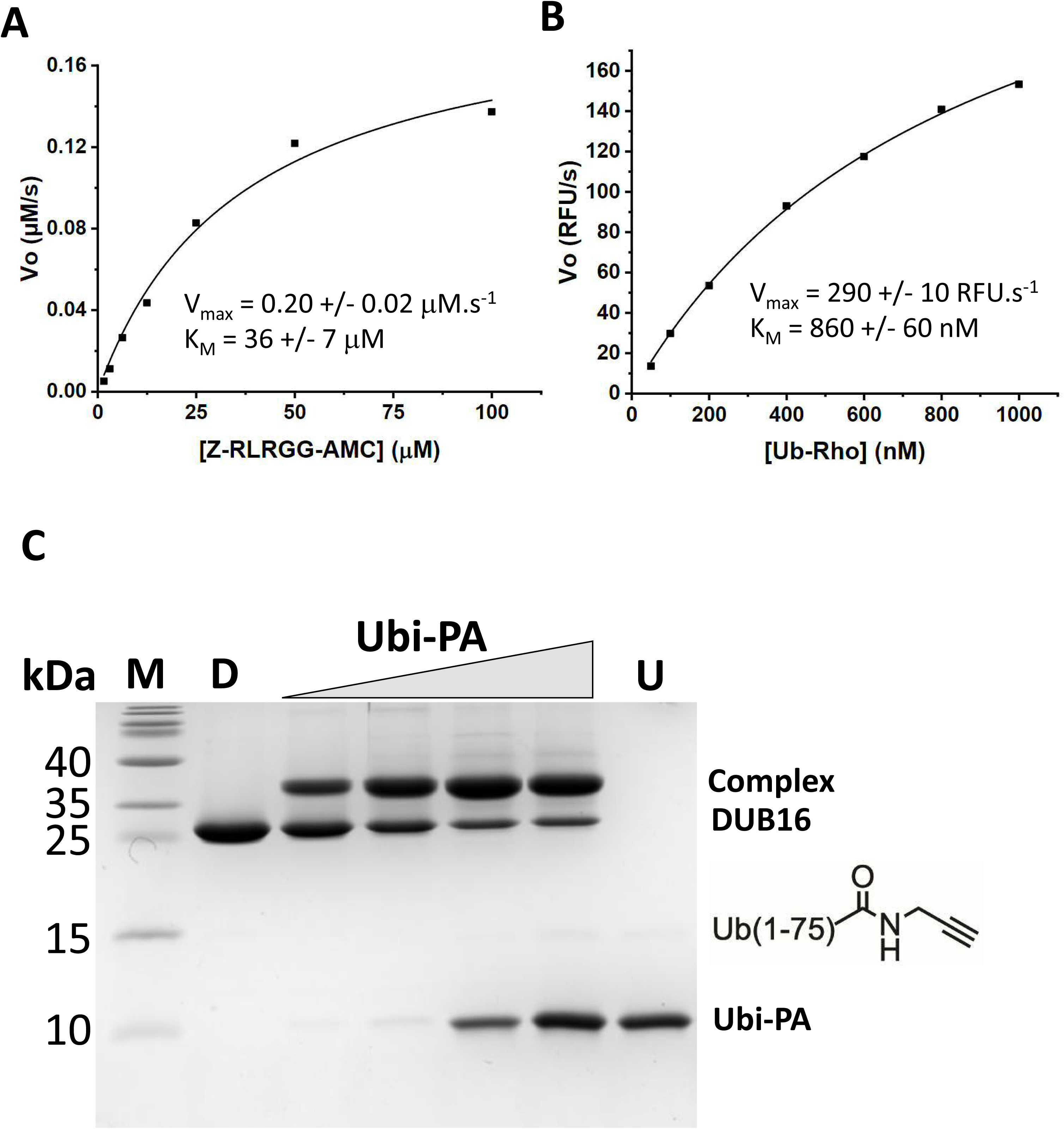
DUB16 acts on Peptide and Ub conjugates. Plots of initial velocity versus substrate concentration for DUB16 cleavage of Z-RLRGG-AMC (**A**) and Ub-RhoGly (**B**). **C**. Coomassie stained SDS-polyacrylamide gel following electrophoresis to resolve DUB16, Ub-propargylamide (Ubi-PA) and their reaction product following mixing and co-incubation with increasing amounts of Ubi-PA to give molar ratios of 0.5, 1, 2, and 4. M – molecular weight markers, D – 3 μg DUB16, U – 2 μg ubiquitin

Next, we measured the activity of DUB16 against full-length ubiquitin covalently linked at its C-terminus to rhodamine 110 Gly (Ub-Rho). The time courses and the fit to the Michaelis-Menten equation are shown in **Figure 1B; Supplementary Figure S1B**, giving a V_max_ of 290 +/− 10 RFU/s and a K_M_ of 860 +/− 60 nM. Assuming that 200 fluorescence units corresponds to 1 nM RhoGly product, this translates to a k_cat_ of 30 s^−1^.

More qualitative assays were performed with full length ubiquitin covalently linked at its C-terminus to 7-amino-4-methylcoumarin (Ub-AMC). In these assays 10 nM DUB16 was incubated with up to 1000 nM substrate (**Supplementary Figure S1C)**. The data show the enzyme is able to turnover ∼100 substrate molecules in 5 minutes, which is comparable to the turnover rates with the shorter Z-RLRGG-AMC substrate. These rates are similar to those measured for the human homologue UCHL1 for the same substrate (32) but approximately 10-fold slower than the rates for human UCHL3 and UCHL3 from *Plasmodium falciparum* (33). Since human UCHL3 and PfUCHL3 also exhibited deneddylation activity (33), we assayed DUB16 against the substrate Nedd8-AMC over the concentration range 100 nM to 2 μM. DUB16 shows activity against Nedd8-AMC (**Supplementary Figure S1D**) demonstrating that the *Leishmania* DUB16 too has dual activity.

Finally, we incubated DUB16 in the presence of the activity-based probe ubiquitin (1–75) propargylamide (Ubi-PA) in which the C-terminal glycine of ubiquitin is replaced with an alkyne containing warhead **(Figure 1C)**. The catalytic cysteines of deubiquitinases react with the terminal alkyne of the modified ubiquitin to form a vinyl sulphide linkage (34) giving rise to products in which the DUB is covalently attached to ubiquitin. After incubation for 30 minutes, the DUB16 reaction products were resolved by SDS polyacrylamide gel electrophoresis. As shown in **Figure 1C**, a lower mobility species is produced with apparent molecular mass of 35 kDa consistent with attachment of ubiquitin (8.6 kDa) to DUB16 (25.5 kDa) to give a fusion protein product with an expected mass of 34.1 kDa. Increasing the Ubi-PA : DUB16 ratio led to only modest increases in the proportion of the enzyme forming the DUB16-Ub complex. We varied the temperature and pH of incubation and explored the presence and absence of different thiol reagents but we were unable to achieve 100 % complex formation. This suggests that a proportion of the protein is not catalytically active.

Taken together, these data show that DUB16 has deubiquitinase activity against small molecule ubiquitin conjugates as expected based on its similarity in sequence to UCH-type DUBs in other species which share these activities (**Supplementary Table S1**).

### Structure of DUB16

The crystal structure of DUB16 was solved by molecular replacement using the human UCHL3 coordinate set (PDBID: 1UCH) (23) as the search model and the structure refined against data extending to 1.69 Å spacing (**Table 1**). There are two molecules, A and B, in the asymmetric unit of the DUB16 crystals. For chain A, residues Met1 – Lys233 can be traced with the exception of residues 127-130, 160-167, 219-223 and for which electron density is missing and these residues are assumed to be disordered. For chain B, the disorder is less extensive with residues 162-166 missing. An ambiguity in the chain tracing resulting from this disorder suggested the possibility that the structure might be a domain-swapped dimer (35) of chains A and B (**Supplementary Figure S3B**) while elsewhere a significant interface (1270 Å^2^) between a pair of crystallographic symmetry related B molecules also suggested the possibility of dimer formation (36). To investigate these possibilities, we measured the solution molecular mass of DUB16 using size-exclusion chromatography and multi-angle laser light scattering (SEC-MALLS). As shown in **Figure 2A**, DUB16 has a weight average molecular mass of 24.8 kDa. This compares with the mass of 25.5 kDa calculated from the sequence and we conclude that the recombinant protein is a monomer.

**Figure 2.**
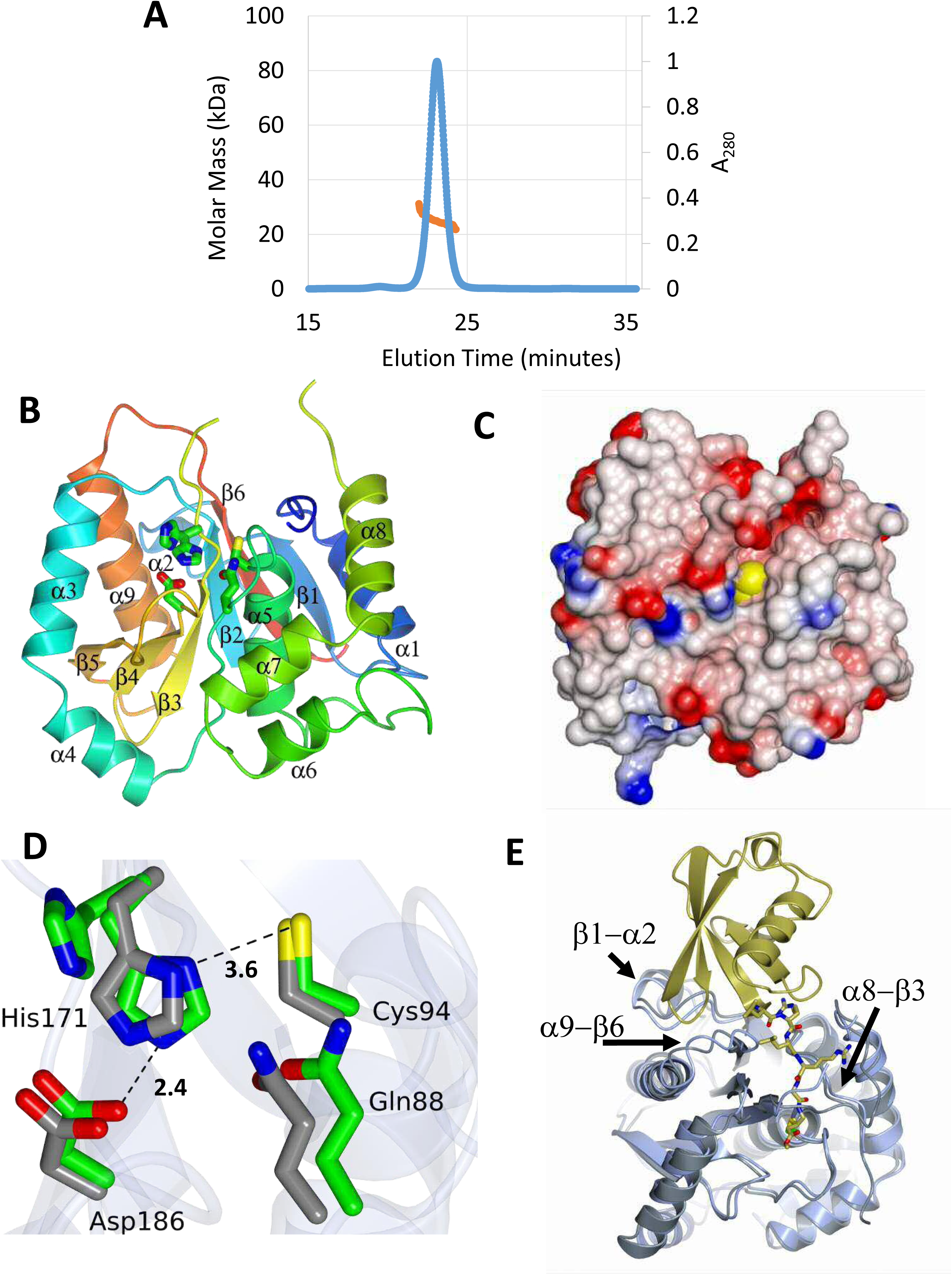
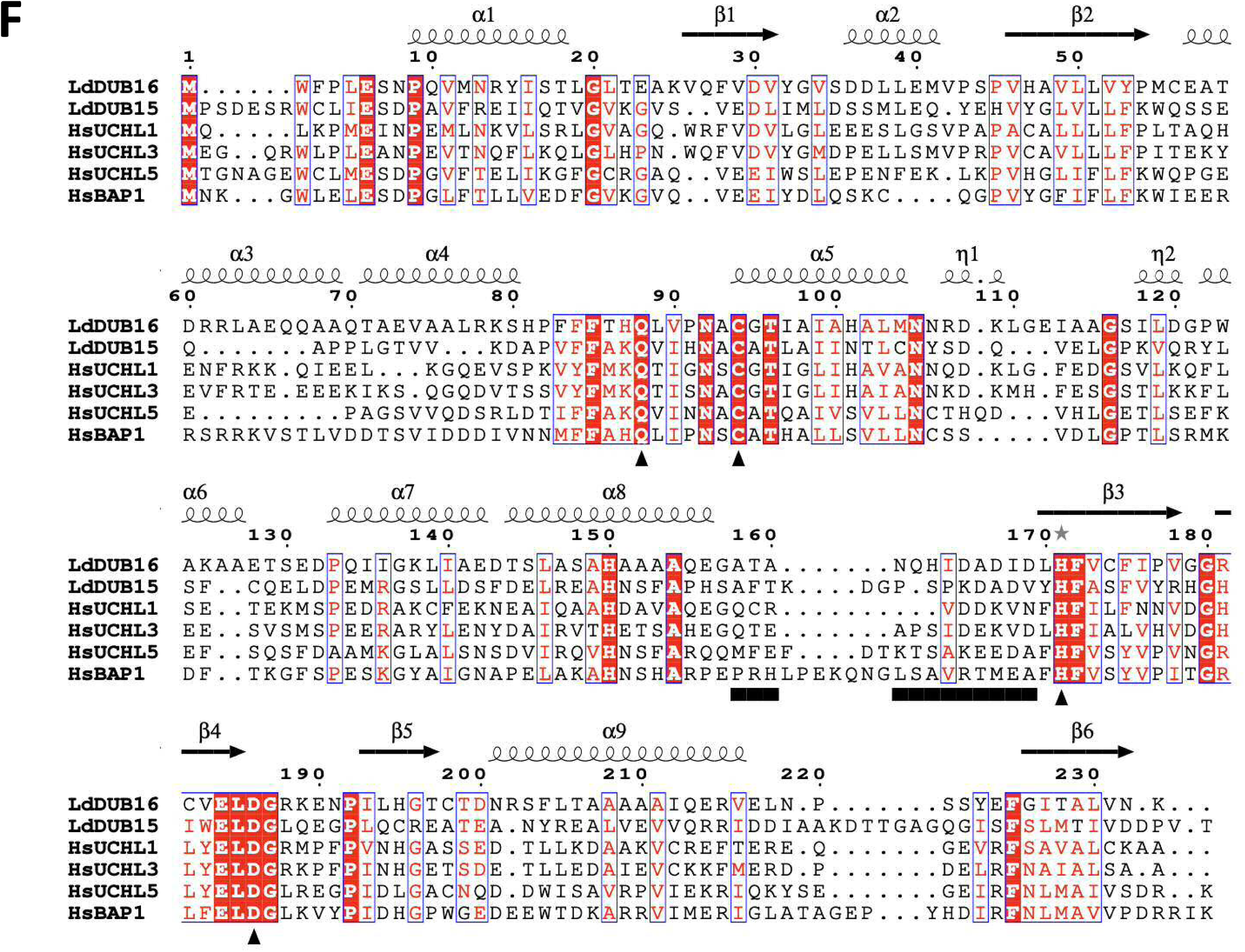
Crystal Structure of DUB16. **A**. Plot of A_280_ (blue line) versus elution time in a SEC-MALLS experiment performed for LdDUB16. The major peak at 23 minutes constitutes >90 % of the sample loaded and is associated with a molecular mass of 25 kDa (red line) calculated from the light scattering and refractive index measurements. **B.** Ribbon tracing of the backbone structure of DUB16. The chain is colour-ramped from the N-terminus (blue) to the C-terminus (red). The secondary structure elements are labelled and the active site residues are shown in cylinder format. **C**. The structure of DUB16 displayed as an electrostatic surface rendering (red, negative, blue positive). The view is essentially orthogonal to that in **B**, looking down into the active site groove. The atoms of the catalytic cysteine 94 residue are shown as yellow spheres emphasising the recessed location of this residue. **D**. The active site residues in the structure of DUB16 superimposed on the active site of the human UCHL3-UbVME complex following superposition of the main chain atoms of the A subunits in the two structures (right). The structures are distinguished by the green and grey carbon atoms respectively. The UbVME moiety has been omitted from the UCHL3 structure for clarity **E**. Superposition of DUB16 (ice blue) and the human UCHL3-UbVME complex (UCHL3, grey: UbVME, gold). The chains are shown as ribbons except for the C-terminal residues of UbVME which are shown as cylinders coloured by atom type with carbons in gold, nitrogens in blue and oxygens in red. The side chains of the active cysteines of the DUBs are shown as cylinders with the carbons in green and the sulphurs in yellow. Elements of the DUB16 secondary structure which partially clash with the UbVME moiety, including the α8-β3 crossover loop are labelled. **F**. Alignment of the sequences of the deubiquitinase domains of the UCH type DUBs prepared in the program TCoffee and displayed in the program ESPRIPT in the context of the secondary structure elements of DUB16. Invariant residues are highlighted by a red background – conserved residues by blue boxes. The black bar below the sequence spans the residues of the crossover loop, the triangles denote the active site residues. The asterisk above the sequence indicates alternate conformations of the residue.

**Table 1.** X-ray diffraction data and refinement statistics.

| **PDB accession code** | **LdoDUB16**<br>**9I77** |
| --- | --- |
| Cell dimensions $a, b, c$ (Å) | 86.40, 86.40, 125.57 |
| Cell angles $\alpha, \beta, \gamma$ (°) | 90.0, 90.0, 120.0 |
| Space Group | $P3_121$ |
| **Data collection** |  |
| Beamline/Wavelength (Å) | DLS i03 / 0.9762 |
| Detector type | CMOS Pilatus 2M |
| Images x oscillation (°) | 3600 x 0.1 |
| Resolution (Å) | 75–1.69 (1.72– |
| $R_{\text{sym}}$ (%)<sup>b</sup> | 6.0 (37.4) |
| $I/\sigma I$ | 19.4 (0.5) |
| Completeness (%) | 100 (99.8) |
| Redundancy | 19.7 (18.3) |
| **Refinement** |  |
| No. unique reflections | 60250 |
| $R_{\text{work}} / R_{\text{free}}$<sup>c</sup> | 21.1 / 24.3 |
| No. atoms | 3647 |
| Protein chain A | 1666 |
| Protein chain B | 1768 |
| water | 213 |
| B-factors (Å<sup>2</sup>) |  |
| all atoms | 44.7 |
| Protein chain A | 46.7 |
| Protein chain B | 42.3 |
| water | 49.1 |
| R.m.s. deviations<sup>d</sup> |  |
| bond lengths (Å) | 0.007 |
| bond angles (°) | 1.619 |

<sup>a</sup> Highest resolution shell is shown in parentheses.
<sup>b</sup> $R_{\text{sym}} = \sum_h \sum_l |I_h - \langle I_h \rangle| / \sum_h \sum_l \langle I_h \rangle$ , where $I_l$ is the $l^{\text{th}}$ observation of reflection $h$ and $\langle I_h \rangle$ is the weighted average intensity for all observations $l$ of reflection $h$ .
<sup>c</sup> $R_{\text{cryst}} = \sum ||F_o| - |F_c|| / \sum |F_o|$ where $F_o$ and $F_c$ are the observed and calculated structure factor amplitudes, respectively. $R_{\text{free}}$ is the $R_{\text{cryst}}$ calculated with 5% of the reflections omitted from refinement.
<sup>d</sup> Root-mean-square deviation of bond lengths or bond angles from ideal geometry.

The A and B chains can be superposed by least squares methods to produce an rmsΔ of 0.7 Å for 217 equivalent Cα atoms. DUB16 comprises a single domain with an α/β fold assembled on a central six-stranded anti-parallel β-pleated sheet (**Figure 2B**). The β-sheet, with a strand order β1-β6-β2-β3-β4-β5, is surrounded by α-helices, with helices α5 and α9 prominent in their close packing onto opposing faces of the sheet. The closest DUB16 structural homologues identified in PDBeFOLD were the human ubiquitin C-terminal hydrolases UCHL1, UCHL3 and UCHL5, and UCHL3 from *P. falciparum*, with Z-scores in the range of 11-14 and rmsΔ values of 1.6-2.0 Å for 180-210 matched C_α_ atoms.

### Active Site

The active site cysteine (Cys94) is located at the N-terminus of the buried helix α5, with its side chain at the base of a groove (**Figure 2 B & C**). From sequence alignments, the other catalytic residues are expected to be His171 and Asp186 which combine with Cys94 to form the catalytic triad, and Gln88 (**Figure 2D & F**). According to the classical cysteine protease mechanism, the imidazole of His171 functions as a base to deprotonate the cysteine thiol, promoting nucleophilic attack at the carbonyl carbon of the scissile peptide (or isopeptide) bond of the substrate. The carboxylate of Asp186 facilitates this process by stabilising the protonated imidazole through the formation of a salt bridge while the side chain amide of Gln88 stabilises the negative charge on the oxyanion that develops in the transition states of the reaction during the formation and breakdown of an acyl enzyme intermediate. In both the A and the B molecules of the structure of DUB16, the side chain of His171 adopts two conformations. In one of these conformers, the imidazole makes a potential ion-pairing interaction with the carboxylate of Asp169 and the cysteine thiol distances are ∼7 Å suggesting that this structure does not represent an active state (**Figure 2D**). This may explain the incomplete complex formation with Ubi-PA (**Figure 1C**). Inactive conformations are common among the structures of ubiquitin C-terminal hydrolase family proteins, and the presence of substrates and/or regulatory components are often required to generate the active states.

We prepared and purified the DUB16:Ubi-PA complex (**Figure 1C**) with a view to determining its structure but were unable to obtain crystals suitable for X-ray analysis. In the absence of a cognate complex structure, we compared the active site of DUB16 with that of the ubiquitin vinyl-methylester (Ub-VME) adduct of human UCHL3 (PDBID 1XD3) (26) following superposition of the deubiquitinase components (**Figure 2E**). Binding of the ubiquitin moiety in DUB16 would require relatively small changes in (i) the conformation of the α9-β6 and α8-β3 loops to accommodate the C-terminal tail of Ub and (ii) the β1-α2 loop so that hydrophobic interactions with the β1-β2 loop of Ub may be formed. The change in the conformation of the α8-β3 loop could presumably take place in concert with the change in the rotamer of the catalytic His171, which resides at the N-terminus of strand β3, to establish the fully active form of the enzyme.

### The Crossover Loop

A much-discussed feature in the structures of UCH family deubiquitinases is the crossover loop. It is so-called because it runs across the top of the active site and appears to restrict access to it (**Figure 2B, E and F**). In DUB16, it links helix α8 to strand β3 and in the A and B molecules of the asymmetric unit, this twelve residue segment is disordered with four and seven residues respectively missing from the electron density maps. In the structure of human UCHL3, a longer 20 residue element of the structure spanning the loop and the preceding α-helix is disordered (23). These residues become ordered in the structure of the complex of this protein with Ub-VME where the crossover loop participates in interactions with the covalently attached Ub analogue (26). The crossover loop adopts a similar conformation and forms similar interactions with Ub in the complex of the yeast homologue, Yuh1, with ubiquitin aldehyde (24). A different situation pertains for PfUCHL3, where the crossover loop is ordered and takes up a similar conformation in both the free and UbVME complexed forms of the enzyme (PDBIDs 2WE6 and 2WDT) (33). Moreover, in the free PfUCHL3 structure the Cys-His-Asp triad is aligned for catalysis. Finally, in this vein, crystal structures of free and Ub-VME complexed human UCHL1 revealed ordered crossover loops but catalytically inactive and active site stereochemistries respectively (25, 32). Collectively, these observations suggest that the crossover loop is flexible and that its conformation is not obligatorily linked to that of the catalytic residues.

### DUB16 exhibits linkage specificity in the cleavage of diubiquitin substrates

We next determined the activity of DUB16 towards a panel of diubiquitin substrates. In these substrates, the C-terminal carboxylate of the distal ubiquitin is joined through an isopeptide bond to one of the seven lysine (Lys6, Lys11, Lys27, Lys29, Lys33, Lys48 and Lys63) side chains of the proximal ubiquitin. In addition, the Gly76 carboxylate can be joined through a peptide bond to the N-terminus (Met1) of the proximal ubiquitin to give linear diubiquitin. Following incubation of the enzyme with the diubiquitin substrates for 30 minutes, the products were resolved by SDS PAGE and visualised by Coomassie staining. As shown in **Figure 3A**, DUB16 cleaved the substrates with linkages at Lys11, Lys48 and Lys63 as evidenced by the disappearance of the diubiquitin bands concomitant with the appearance of higher mobility ubiquitin bands. There was very low or no activity towards linear diubiquitin or substrates with the Lys6, Lys27, Lys29 and Lys33 linkages. Even for the Lys11, Lys48 and Lys63 substrates, at the close to equimolar enzyme to diubiquitin ratios used, the turnover is very low. While the disappearance of the diubiquitin species is clear, the monoubiquitin product bands are diffuse. We therefore subsequently used Western blotting and detection with an anti-ubiquitin antibody to visualise the ubiquitin-containing products (**Figure 3B**). In these experiments, we included a 5-minute time point. This gel emphasises the linkage selective action of DUB16 on the diubiquitin substrates with activity increasing in the order Lys63 ≤ Lys48 < Lys11.

**Figure 3.**
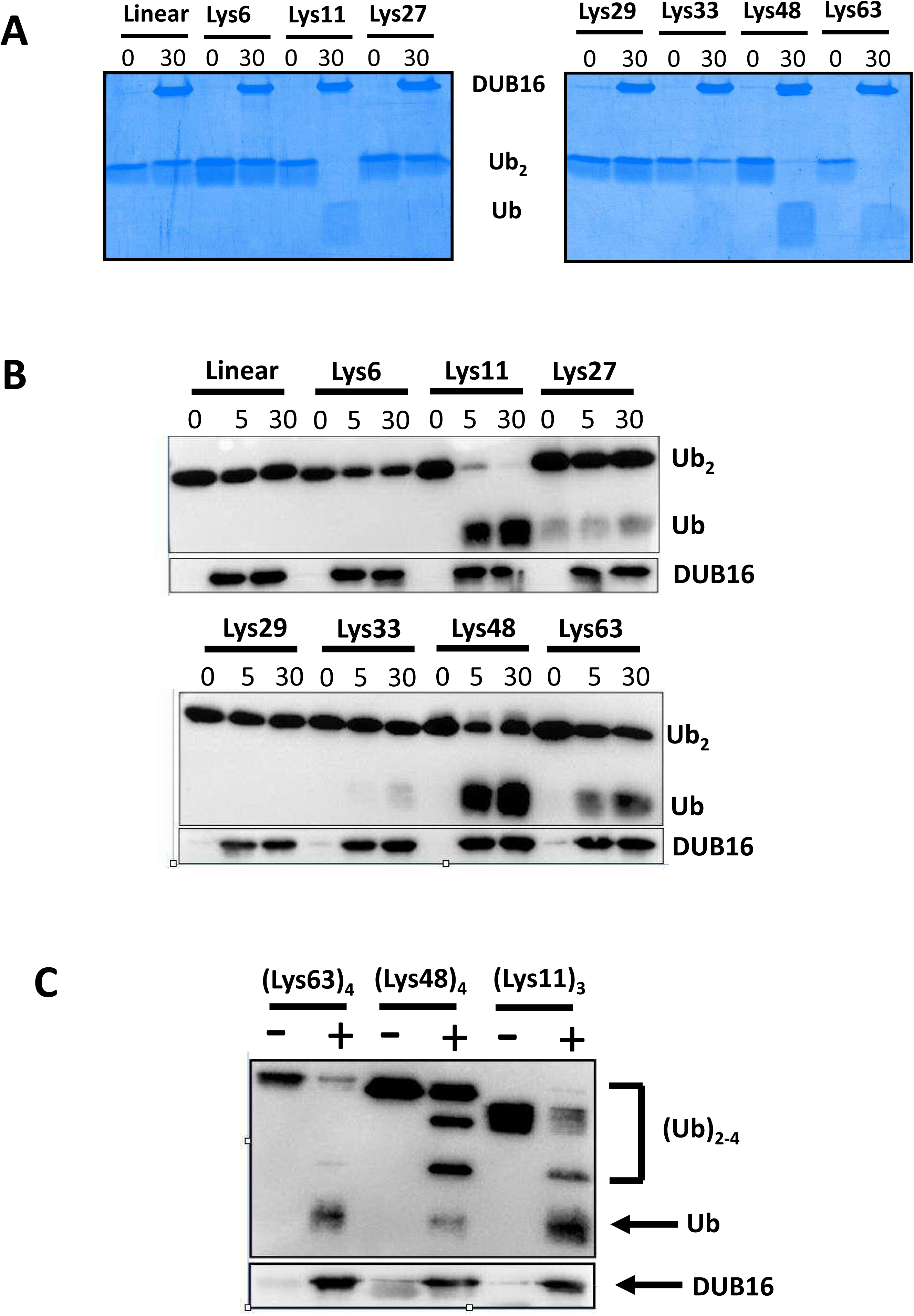
DUB16 displays linkage specificity in the cleavage of diubiquitin substrates. Deubiquitinase and diubiquitin substrates with the indicated linkages were co-incubated for the indicated time in minutes prior to quenching with the addition of SDS-containing sample buffer and resolution of the products by 12 % polyacrylamide gel electrophoresis. In **A**, the gel was stained with Coomassie blue. In **B**, following Western transfer and probing with anti-ubiquitin antibody, the blots were visualised with the ChemiDoc system (BioRad). Mouse anti-DUB16 antibody (51) was used to immunoblot the enzyme as a loading control. **C**. Cleavage of tri- and tetra-ubiquitins monitored as in B.

Next, DUB16 activity was assayed against longer ubiquitin chains, specifically Lys11-linked triubiquitin, and Lys48- and Lys63-linked tetraubiquitins. The reaction products formed after a 30-minute incubation were resolved by SDS-PAGE and probed following Western blotting with the anti-Ub antibody (**Figure 3C**). For the Lys48-tetraubiquitin substrate tri-, di- and monoubiquitin species were present among the products while for the Lys11-linked triubiquitin substrate, diubiquitin and monoubiquitin species were observed. Meanwhile for the Lys63-tetraubiquitin substrate, triubiquitin species were not observed, with a faint band for the diubiquitin product observed together with monoubiquitin. The absence of a triubiquitin species may indicate preferential cleavage of the central linkage to produce a pair of diubiquitins. In all three experiments, some uncleaved substrate was observed.

### DUB16 cleaves a linear fusion of Ub to ribosomal protein L40

In *Leishmania* species, ubiquitin is encoded either as chains comprising 30 or more ubiquitin repeats linked in a linear head-to-tail manner, or as an N-terminal fusion to the ribosomal protein L40 (**Figure 4A**). There are typically three copies of the gene encoding the Ub-L40 fusion. The AlphaFold2 structure of Ub-L40 shows the linker segment Arg-Gly-Gly-Val-Met in an extended conformation (**Figure 4A**). Production of the mature ubiquitin and L40 proteins requires proteolytic cleavage of the Gly-Val peptide bond. For activity assays, we amplified the LdBPK_311930 Ub-L40 coding sequence (**Supplementary Table 2**) and overproduced and purified the recombinant protein, with an N-terminal Met-Ala and a C-terminal Leu-Glu-His-His-His-His-His-His purification tag. Mass spectrometry analysis of the purified product revealed a prominent species with a mass of 15,816.4 Da. This is a close match to that expected for the recombinant Ub-L40 protein lacking the N-terminal methionine (15,818.4 Da). We attribute the remaining 2 Da difference to a disulphide bridge forming on the L40 component (**Supplementary Figure 4**).

**Figure 4.**
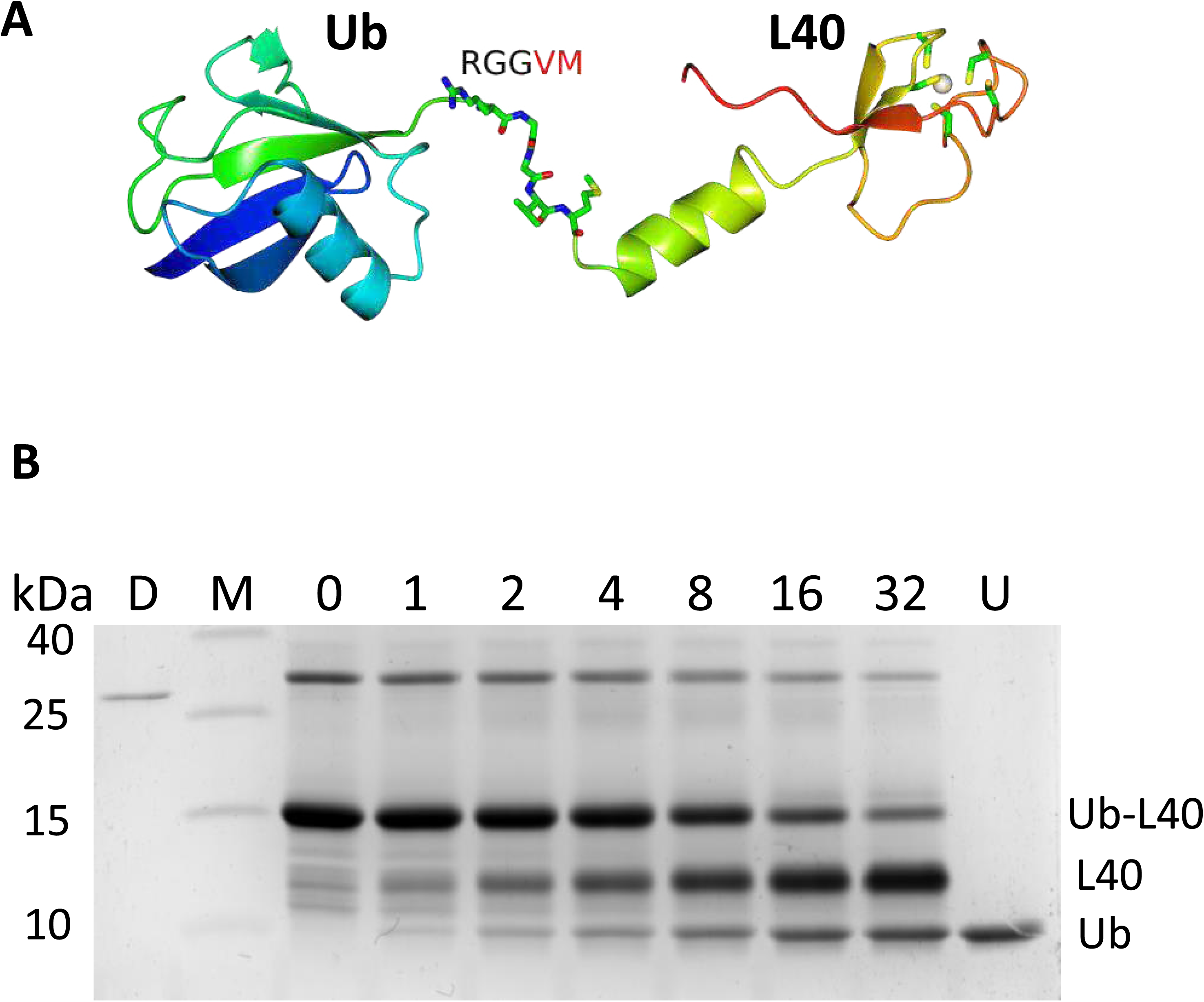
DUB16 cleaves the Ubiquitin-L40 precursor protein to produce monomeric ubiquitin. **A**. Ribbon rendering of the AlphaFold2 model AF_E9BMW6. A zinc metal is shown as a white sphere, with its surrounding cysteines and residues in the linker region shown as cylinders. **B**. Time course of Ub-L40 cleavage by DUB16. 10 μg of Ub-L40 substrate was incubated with 30 ng of DUB16 at 24°C and reaction aliquots were removed over the time range 0 - 32 minutes and quenched in SDS PAGE sample buffer. The reaction products were resolved by denaturing 17.5 % polyacrylamide gel electrophoresis and visualised by staining with Coomassie blue dye. The disappearance of the Ub-L40 band of apparent Mr = 16 kDa is accompanied by the concomitant appearance of the two higher mobility species, L40 and Ub. D = DUB16 (300 ng), M = markers, U = Ubiquitin (3.5 μg).

Purified Ub-L40 was incubated in the presence of DUB16 and the reaction products were analysed by SDS polyacrylamide gel electrophoresis followed by staining with Coomassie blue dye. As shown in **Figure 4B**, Ub-L40 (136 residues, 15.8 kDa) migrates with similar mobility to the 15 kDa molecular weight marker. Following treatment with DUB16, two higher mobility species appear which we attribute to the L40 and Ub products. The mobility of the latter matches that of a control ubiquitin sample, while Western blotting of similar samples probed with anti-ubiquitin antibodies confirmed the presence of ubiquitin in this faster migrating band (**Supplementary Figure 2**). The L40 protein has an Arg/Lys rich sequence and a theoretical pI of 10.3, which may account for its anomalously low electrophoretic mobility in denaturing polyacrylamide gels. Mass spectrometry analysis of the reaction products produced a peak at 8619.4 Da which is an exact match to the mass of the ubiquitin product, assuming that cleavage has taken place after Gly76 (**Supplementary Figure 4**). The C-terminal L40 product was not observed in the mass spectrum.

Judging from the extent of staining in these gels, half of the starting material (∼600 pmol) has been cleaved in ∼10 minutes by ∼1 pmol enzyme corresponding to a turnover of ∼0.5 per second. This is lower than that for the cleavage rate of the Ub-small molecule conjugates but considerably higher (10^3^-10^4^-fold) than that for cleavage of the diubiquitins. This is a functionally interesting result as it shows that DUB16 is able to cleave the peptide bond between Ub and L40 but not that linking the ubiquitins in linear diUb. In the crystal structure of linear diubiquitin (PDB Entry: 2W9N) (30), a short unstructured segment of the polypetide links the two discrete Ub domains. This unstructured region is shorter than that linking the Ub and L40 elements in the AlphaFold model of Ub-L40, where the scissile peptide bond appears to be much more accessible to proteolytic cleavage by DUB16.

### Cleavage of Ub-L40 is inhibited by cyanopyrrolidines IMP-1711 and IMP-1710

A potent and selective cyanopyrrolidine inhibitor of human UCHL1 (IC_50_ = 90nM), (S)-2-(4-(1H-pyrrolo[2,3- b]pyridin-3-yl)indoline-1-carbonyl)pyrrolidine-1-carbonitrile (IMP-1711 in **Figure 5A),** with cellular activity has been reported (37). In view of the sequence similarity shared by DUB16 and UCHL1 (**Figure 2F, Supplementary Table 1**), we tested whether IMP-1711 could inhibit the *Leishmania* enzyme. IMP-1711 is a cysteine-reactive covalent DUB inhibitor which is expected to react with the thiol of Cys94. As shown in **Figure 5B**, in the presence of excess IMP-1711, there is little or no cleavage of the Ub-L40 in the gel-based assay. Next, we assessed the binding of IMP-1711 to DUB16 using differential scanning fluorimetry. As shown in **Figure 5C**, in the presence of a 2- fold excess of IMP-1711, the melting temperature of DUB16 increased by ∼7 °C as judged by the change in fluorescence of the SYPRO Orange dye which is expected to bind to the unfolded form of the protein. The significant change in T_m_ is consistent with covalent binding of IMP- 1711 to DUB16.

**Figure 5.**
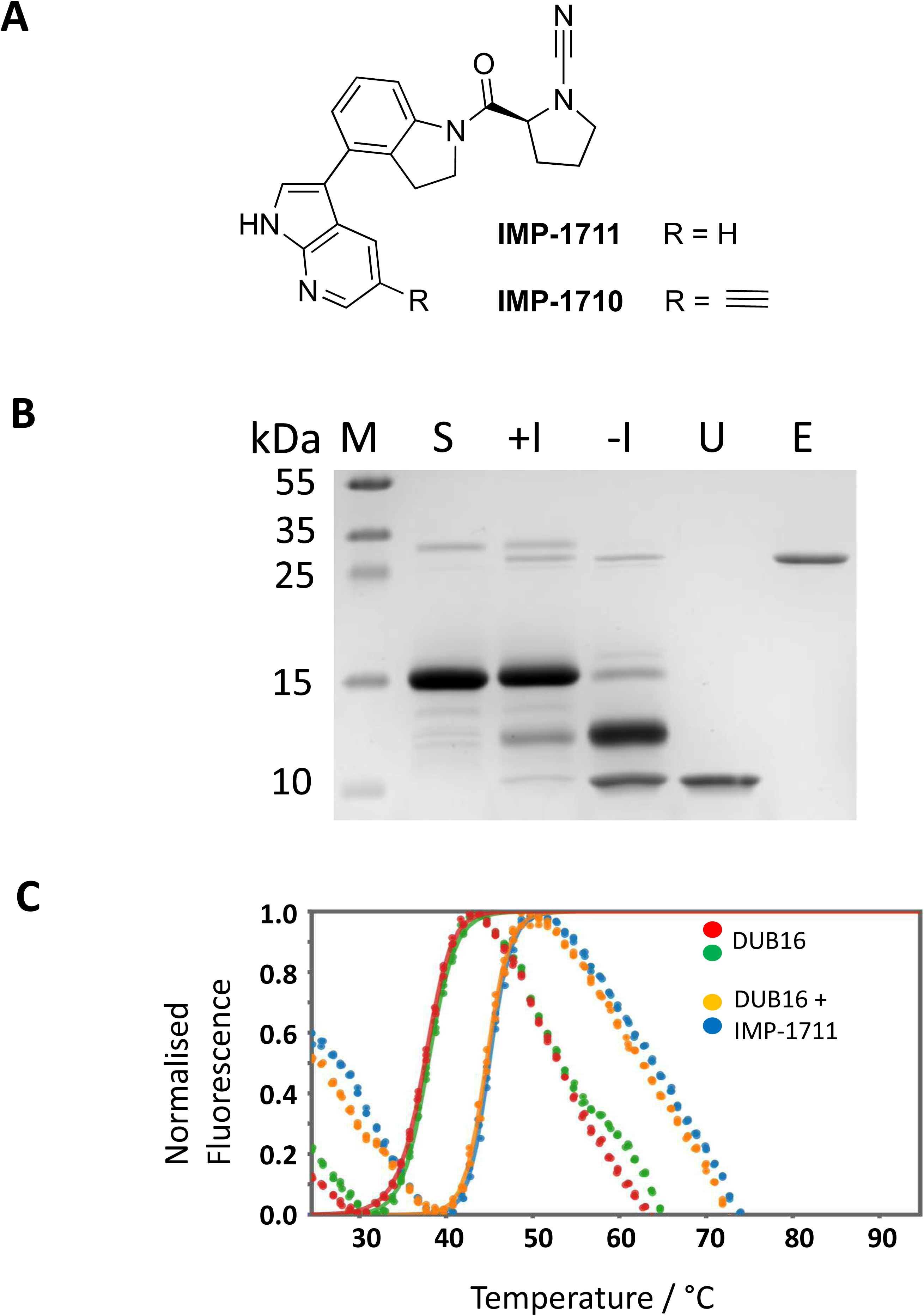
*In vitro* inhibition of DUB16 activity. **A.** Structures of the cyanopyrrolidine IMP-1711 and its derivative IMP-1710 bearing an alkyne tag. **B**. Coomassie stained SDS-PAGE showing inhibition of Ub-40 cleavage by IMP-1710; M = Mwt markers, S = Ub-L40 substrate, +/-I corresponds to DUB16 preincubated (30 min) in the presence and absence of a 40-fold excess of IMP-1710 before addition to the Ub-L40 substrate and further incubation at room temperature for 30 min, U = 2 μg ubiquitin, E = 0.5 μg DUB16. **C**. Differential scanning fluorimetry profile of 12.5 μM DUB16 in the presence and absence of 25 μM IMP-1711.

### Anti-Leishmanial activity of IMP-1710 and target deconvolution

Following the observation that IMP-1711 can inhibit DUB16 *in vitro*, we explored whether it had anti-leishmanial activity against *L. donovani* LV9. A cell viability assay was conducted after exposing promastigotes to varying concentrations of IMP-1711, revealing it had an EC_50_ of 0.76 +/- 0.12 μM (**Table 2**) – for comparison, the EC_50_ of miltefosine against this strain in our hands is ∼1.5 μM (38).

**Table 2:** Results of dose-response cell viability assays. The table lists the cell line, the compounds tested and the mean EC_50_ (with 95% confidence intervals reports). The mean is derived from 3 experiments conducted each with 3 technical replicates (N=9).

| Cell Line | Compound | EC50 μM (95% CI) |
| --- | --- | --- |
| *L. donovani* LV9 | IMP-1711 | 0.76<br>(0.64 - 0.88) |
| *L. mexicana* M379 | IMP-1710 | 2.17<br>(2.03 - 2.33) |
| *L. mexicana* M379 [DUB16]<br>Population A | IMP-1710 | 3.65<br>(3.45 - 3.84) |
| *L. mexicana* M379 [DUB16]<br>Population B | IMP-1710 | 3.21<br>(2.94 - 3.49) |

To determine whether the anti-leishmanial effect of UCHL1 inhibitors was due to activity against DUB16, we elected to test if overexpressing DUB16 would confer increased resistance to the treatment. Here we switched to *L. mexicana*, a genetically more tractable species that causes cutaneous leishmaniasis. *L. mexicana* M379 promastigote forms were transfected with a pNUS episome (10, 39) to generate two independent populations that overexpress an untagged form of DUB16, (*L. mexicana M379* [*DUB16*]). To establish the relative amounts of active DUB16 in each strain, an activity-based protein profile was performed using an HA-Ubiquitin-propargylamide probe (HA-Ubi-PA) (10). HA-Ubi-PA is recognised by a broad spectrum of deubiquitinase enzymes and reaction of the propargyl group with the active site cysteine irreversibly inhibits cysteine protease DUBs by forming a vinyl thioether linkage. The HA epitope allows for enrichment and/or detection of the bound DUBs using anti-HA antibodies.

Cell lysates prepared from the parental strain and each of the [*DUB16*] strains were incubated with HA-Ubi-PA, the reaction was then terminated by the addition of lithium dodecyl sulfate sample buffer (LDS). The protein samples were resolved by SDS-PAGE then analysed by Western blotting against the HA epitope. An HRP-conjugated secondary antibody allowed for chemiluminescence determination of the amount of DUB activity in the sample. A prominent band was observed in the [*DUB16*] populations with an apparent molecular weight of ∼35 kDa, which would correspond to the mass of DUB16 (25.5 kDa) plus the mass of HA-Ubi-PA probe (9.9 kDa); this was consistent with the signal observed by Damianou (2020). The chemiluminescent signal intensity of this band was quantified using ImageLab software, suggesting that DUB16 activity increased 90-300-fold in the two independent parasite populations containing the episome (**Figure 6A & B**).

**Figure 6.**
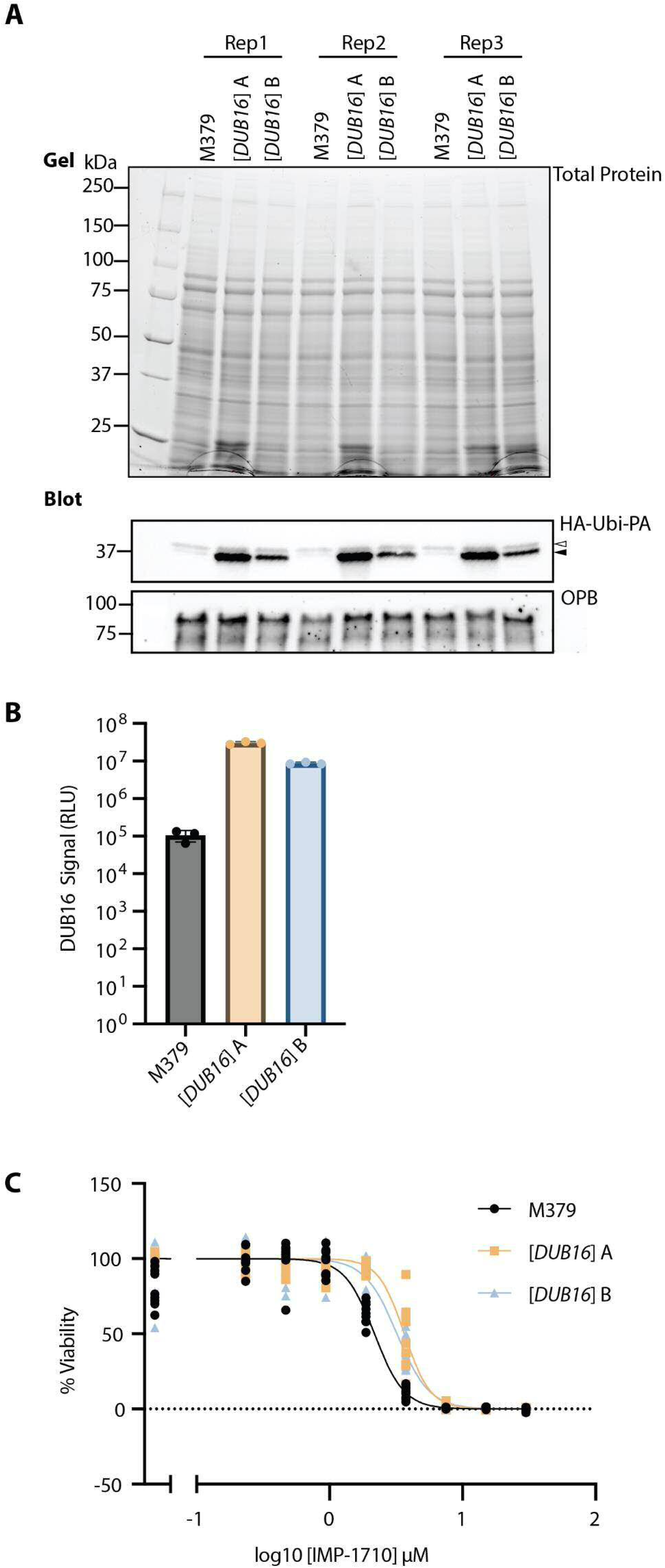
*In vivo* profiling of inhibitor activity. **A.** DUB activity-based probe assay. Two populations (labelled as [*DUB16*] A & B) of *L. mexicana* M379 promastigotes were transfected with the *pNUS::DUB16* episomal expression vector and selected with G418. Cell lysates were incubated with the HA-Ubi-PA activity-based probe and resolved by SDS-PAGE. Total protein in the polyacrylamide gel was visualised with the BioRad Stain Free TGX system prior to transfer to a PVDF membrane. Western blotting with an anti-HA primary antibody and visualising with an HRP secondary antibody shows upregulation of DUB16 activity in the transfected populations. In the HA-Ubi-PA blot, the lower band (indicated by the solid arrowhead) represents DUB16, the upper band (indicated by the open arrowhead) is likely to be DUB17 (10). Oligopeptidase B (OPB) is used as a loading control for the blot. **B.** Bar chart representing the mean anti-HA signal corresponding to DUB16 activity, quantified using BioRad ImageLab. Error bars denote the standard deviation. **C**. Does response curves showing the activity of IMP-1710 against promastigote forms of *L. mexicana* M379 and the derived strains overexpressing DUB16 using a cell viability assay. The EC_50_ values are given in Table 2.

Following the validation that DUB16 activity was increased in the [*DUB16*] strains, the two populations were compared to the parental strain using the promastigote cell viability assay (38). The cells were exposed to varying concentrations of IMP-1710, a commercially available derivative of IMP-1711 modified to contain an alkyne group to enable click chemistry, in the expectation that it retains similar bioactivity (37). The EC_50_ of IMP-1710 against *L. mexicana* M379 was 2.17 μM. This rose to 3.65 μM and 3.21 μM in the two independent populations of *L. mexicana M379* [*DUB16*] (**Figure 6C**, **Table 2**). Although modest, the strains overexpressing DUB16 were consistently more resistant to IMP-1710 treatment, suggesting that DUB16 can be targeted in living parasites by this molecule.

## Discussion

Here we have purified recombinant *L. donovani* DUB16 profiled its activity against a range of ubiquitin substrates. In addition, we determined its three dimensional structure revealing a typical ubiquitin C-terminal hydrolase fold in which the catalytic residues reside at the bottom of a prominent substrate-binding groove which is bridged by a crossover loop. DUB16 was shown to cleave small molecule conjugates of ubiquitin and the ubiquitin-like protein, Nedd8. These results are consistent with those obtained for human UCHL1 and UCHL3 which rapidly cleave a range of small molecule Ub conjugates with little discrimination towards the leaving group (27). DUB16 was also shown to cleave diubiquitins with a specific subset of isopeptide linkages, albeit with much lower turnover numbers. It showed no activity towards linear diubiquitin similarly to both UCHL1 and UCHL3 (27). Finally, DUB16 was able to cleave ubiquitin fused to the ribosomal protein L40 with a turnover number similar to that of UCHL3 and considerably faster than that of UCHL1 (27).

The faster rates of hydrolysis of small molecule *versus* larger protein conjugates seen here and reported previously for UCH family DUBs has been attributed to the crossover loop which restricts access to the active site. In one scenario, the crossover loop is a barrier to ligand binding. For the ubiquitinated substrate to engage productively with the enzyme, the C-terminally conjugated moiety has to be threaded through the narrow space between the crossover loop and the catalytic site until the carbonyl group of Gly76 of Ub is appropriately juxtaposed with the active cysteine. This would restrict activity to smaller substrates such as the synthetic aminocoumarin adducts for which DUB16 exhibits deubiquitinase and deneddylase activities, and short peptide adducts of ubiquitin which might otherwise accumulate *in vivo* as a result of cellular processing. The capacity of the DUB domains of UCHL1, UCHL3, UCHL5, and BAP1 to cleave K48-linked diUb was investigated by engineering deletions and insertions into the crossover loops (40). This study concluded that crossover loop lengths greater than 14 residues are required for cleavage of the diubiquitin, with shorter loop lengths limiting cleavage to smaller Ub-conjugates.

In a second scenario, the crossover loop can peel away to allow access of larger targets to the active site. According to this model, the crossover loop adopts an ‘out’ conformation allowing substrate binding and a locked ‘in’ conformation which stabilises the acyl enzyme upon release of the leaving group (26). Consistent with the structural data, in the absence of substrate, the crossover loop can adopt either or neither of these conformations, while in the acyl-enzyme complex it will have the locked ‘in’ conformation.

### Linkage specificity

The data presented here indicate that DUB16 has linkage-specific diubiquitin cleavage activity preferring substrates with Lys11, Lys48 and Lys63 linkages. This set of linkage preferences is reminiscent of the deubiquitinase Cezanne (16) although turnover numbers are much higher for the latter and the Lys11 linkage preference is even stronger. While it has yet to be confirmed in *Leishmania*, Lys48-linkages are elsewhere the most abundant linkage and associated with proteasomal degradation (12). Lys11 linkages are also associated with proteasomal targeting particularly in the context of cell cycle regulation while Lys63 linkages are associated with cell signalling, trafficking and DNA repair (12). Experimental data on the abundance and types of ubiquitin linkages present in *Leishmania* parasites are very limited (9). It was shown that Lys63-linked diubiquitin can be generated by UBC2-UEV1 from *L .mexicana* (13) while at close to stoichiometric ratios the deubiquitinase DUB2 can cleave all diubiquitin linkage types except Lys27 (10).

### De novo ubiquitin synthesis and essential function of DUB16

In eukaryotic cells, deubiquitinases play important roles in *de novo* ubiquitin synthesis. This is because ubiquitin is synthesised in precursor forms that must be processed to yield the active molecule. In many organisms, four genes encode ubiquitin precursors. These comprise single ubiquitin molecules C-terminally linked to either ribosomal protein L40 or ribosomal protein S31, and polyubiquitin forms comprising numerous (up to 10 in mammals) head-to-tail linked ubiquitin units. These polyubiquitin chains invariably possess an amino acid or oligopeptide extension that caps the C-terminal glycine ensuring that the ubiquitin precursor is not inappropriately conjugated to cellular proteins (27, 41). It is likely that cleavage of these C-terminal entities would be catalysed by DUB16 amongst other deubiquitinases. In *Leishmania* species, three genes encode ubiquitin fused to ribosomal protein L40, with a fourth gene encoding polyubiquitin. The polyubiquitins of these parasites are exceptionally large (18) with the LMJFC_360049300 gene of *L. major* strain Friedlin 2012 encoding 75 tandem ubiquitin units (https://tritrypdb.org/) (42).

DUB16 is the smallest of the 20 cysteine protease DUBs found in *Leishmania* with its crystal structure exhibiting a single compact domain. It is one of only four DUBs that are essential for the viability of promastigote stage parasites (10). Very little is known of the repertoire of ubiquitinated proteins, or the dynamics of ubiquitination in *Leishmania* and how it is shaped by DUBs which are likely to have overlapping substrate ranges. As a result, speculation on the essential function(s) of DUB16 is difficult. Nevertheless, its activity towards Ub-L40 warrants consideration since both ubiquitin and the ribosomal L40 subunit are likely to be essential in the parasite. Since ubiquitin is additionally available through cleavage of polyubiquitin chains, it seems less likely that the parasite depends on DUB16 to replenish the ubiquitin pool from this source. Moreover, since this enzyme does not cleave the linear linkage in diubiquitin, it is unlikely to process polyubiquitin indicating that this precursor is processed by other enzymes.

In contrast, all three of the ribosomal L40 genes encode the ribosomal subunit as a fusion to ubiquitin. In *Saccharomyces cerevisiae*, mutants blocked in ubiquitin-L40 processing are unable to grow (43). The uncleaved precursor assembles into pre-60S ribosomal particles but these are not competent for translation. This effect can be attributed to steric interference by the ubiquitin moiety of elongation factor EF2 binding to the GTPase site on the ribosome (41). Since DUB16 cleaves Ub-L40 efficiently, and Ub-L40 is the only source of the L40 subunit in *L. donovani* and *L. mexicana*, catalysis of this reaction may be an essential function of the enzyme.

To investigate this possibility, we used the cyanopyrroline, IMP-1711 and its derivative IMP-1710, discovered as potent covalent inhibitors of UCHL1 (37). We found that IMP-1711 binds tightly to DUB16 as evidenced by thermal shift analysis and efficiently inhibits the cleavage of Ub-L40 *in vitro*. Moreover, IMP-1710 has cell killing activity against *L. donovani* and *L. mexicana* promastigotes with a potency similar to miltefosine in our assays. To explore whether the cell killing effect of IMP-1710 is due to its *in vivo* targeting of DUB16, we generated *L. mexicana* parasites overproducing the enzyme, reasoning that this would lessen the inhibitor’s potency in cell killing. Indeed, we observed a small but significant increase in EC_50_. Nevertheless, this effect is modest given the high levels of enzyme overproduction achieved. It is likely therefore, that as observed in mammalian cells, more than one deubiquitinase contributes to Ub-L40 processing (44). It is also possible that there are additional intracellular targets of the inhibitor.

### DUBs as therapeutic targets

As dysregulation of deubiquitinases is associated with neurodegenerative disorders, cancer metastasis, and inflammation and immunity in humans, these enzymes have become attractive therapeutic targets (20, 45). Many DUB inhibitors are in early stages of drug development, a few have entered clinical trials though none are currently approved for clinical use. A major hurdle is the similarities among DUB active sites making it difficult to target a single human DUB without off-target effects (20, 46). A legacy of this research effort is a rich catalogue of small molecule deubiquitinase inhibitors and activity-based probes which can advance the validation of DUBs as drug targets for the treatment of infectious parasitic diseases. These reagents enabled profiling of the deubiquitinases of *Leishmania* and advancement of DUB16 as a promising candidate for inhibitor development in leishmaniasis (10). The further aspiration is that some of these existing inhibitors can be repurposed and developed as future anti-leishmanial therapeutics.

## Methods

### Cloning, overexpression and purification of DUB16

A 728 bp DNA fragment spanning the coding sequence of DUB16 (Gene ID: LdBPK_250190.1) was amplified from *L. donovani* BPK chromosomal DNA by PCR using the primer pair DUB16_F and DUB16_R (**Supplementary Table 2**) and Hi-Fi DNA polymerase. The purified PCR product was digested with the restriction endonucleases *Bam*HI and *Hin*dIII and mixed with similarly cut pET28a vector DNA. After incubation in the presence of T4 DNA ligase and ATP, the reaction products were added to competent *E. coli* DH5α cells and following plating, kanamycin-resistant colonies were selected overnight. Transformants were screened by colony PCR, restriction endonuclease digestion of purified plasmid DNA and finally DNA sequencing. The resultant plasmid, pET28a-DUB16, encodes resides 1-233 of DUB16, N-terminally fused to a 34-residue vector-derived sequence containing a thrombin-cleavable hexahistidine tag.

For overproduction of recombinant enzyme, pET28a-DUB16 was introduced into *E. coli* Rosetta (DE3) cells and overnight cultures grown in Luria Bertani (LB) kanamycin medium at 37°C. The following morning, the cells were diluted 100-fold into fresh medium and grown with shaking at 200 rpm until an OD_595_ of 0.4–0.6 was achieved, when IPTG was added to 1 mM to induce expression. After a further 4 hours, the cells were harvested by centrifugation and the pellets stored at −80°C.

For protein purification, the bacterial cells were thawed on ice and resuspended in 10 ml per litre of cell culture of Buffer A (50 mM Tris–HCl, pH 8.0, 300 mM NaCl, 10 mM imidazole, and 10% glycerol) containing 1 mg.ml^-1^ lysozyme. After 30 minutes incubation on ice, cells were sonicated (Probe diameter = 3mm, Misonix, Farmingdale, NY, USA) in 10 cycles of 10 s pulses at 25 W with 1 minute intervals between pulses. The supernatant was recovered following centrifugation of the lysate at 10,000 g for 30 minutes at 4°C and mixed with Ni-NTA agarose resin (1 ml) equilibrated in Buffer A. Following a 1-2 hour incubation at 4°C with moderate agitation, the resin was allowed to settle and poured into a column. The column was washed with 5 volumes of Buffer A and bound proteins eluted in Buffer A containing 240 mM imidazole. The eluting fractions were analysed by SDS polyacrylamide gel electrophoresis and those enriched in a protein of ∼25 kDa molecular mass were pooled and dialysed against (20mM Tris pH 8.0, 150mM NaCl buffer). The yield of protein was estimated using a Bradford assay to be 40-50 mg per litre of culture.

### Preparation of Ub-L40 fusion protein

For the purposes of DUB16 assays, we amplified the ubiquitin-L40 (LdBPK_311930.1) coding sequence from *L. donovani* BPK template DNA by PCR using the primer pair UbL40-F and UbL40-R (**Supplementary Table S2**), and ligated the resulting fragment following *Nco*I/*Xho*I restriction enzyme digestion into similarly cut plasmid pET28a. The resulting plasmid was introduced into the *E. coli* expression strain Rosetta (DE3). A recombinant fusion protein comprising, 76 residues of Ub, 52 residues of L40 and an octapeptide purification tag, LEHHHHHH, was produced at high levels following induction with 0.5 mM IPTG and was purified by Ni^2+^-chelation and size exclusion chromatography to >95 % homogeneity as judged by Coomassie staining of SDS-polacrylamide gels.

### Deubiquitinase activity against Z-RLRGG-AMC

Z-RLRGG-AMC is a substrate in which the C-terminal pentapeptide from ubiquitin is conjugated to 7-amino-4-methylcoumarin (AMC). Cleavage of the substrate produces free aminocoumarin and enhanced fluorescence emission. Z-RLRGG-AMC (Sigma-Aldrich) was serially diluted in reaction buffer (50 mM Tris pH 8, 50 mM NaCl, 5 mM DTT and 0.002 % Tween-20) to concentrations in the range 200-3.125 μM. DUB16 was diluted in the same reaction buffer to a concentration of 200 nM. 30 μl of each Z-RLRGG-AMC dilution was added in triplicate to reaction mixes in a black 96-well plate (Thermo Scientific Nunc). The reactions were initiated by addition of 30 μl DUB16. Reactions were performed at 25°C and readings were taken every 37 seconds for 37 minutes (λ_ex_ = 350-15 nm, λ_em_ = 440-20 nm) with a CLARIOstar plate reader (BMG LABTECH). An AMC standard curve was generated, by measuring fluorescence as a function of AMC (Fluorochem) concentration. The first 10 % of the data points were used to calculate the initial reaction velocity at each substrate concentration, and the initial velocities were plotted against substrate concentration. Data analysis was performed with OriginPro 2023b software. Data were fitted using non-linear curve fit, Michaelis Menten, to determine V_max_ and K_M_.

### Deubiquitinase activity against Ub-Rho110Gly

The format of this assay was similar to that described above. A stock solution of Ub-Rho110Gly obtained from Ubi-Q was diluted to concentrations in the range 2000-100 nM. DUB16 was diluted to 100 pM. 40 μl of each Ub- Rho110Gly dilution was added to wells in triplicate, and the reactions initiated by addition of 40 μl of DUB16. Readings were taken every 40 seconds for 30 minutes (λ_ex_ = 487-514 nm, λ_em_ = 535-30 nm).

### Deubiquitinase activity against Ub-AMC Conjugates

Deubiquitination activity was also measured in a fluorometric assay with the substrate Ub-AMC. 10 nM DUB16 was incubated with varying concentrations of Ub-AMC at 37°C in 2 x DUB buffer (100 mM NaCl, 100 mM Tris, pH 7.4 and 10 mM DTT). The fluorescence signal was recorded over 36 minutes following substrate addition. Since many cysteine proteases possess dual DUB/deNeddylating activity, we repeated this assay using Nedd8-AMC as substrate.

### Cleavage of di- and oligo- ubiquitin substrates

Purified recombinant DUB16 was incubated with a panel of di- or oligo-ubiquitins (Boston Biochem now R&D Systems) containing different Ub-Ub linkages. Reaction mixes containing 1 μg of the ubiquitin substrate in 25 mM Tris, pH 7.5 and 4.5 μM of DUB16 were incubated for up to 45 minutes at 37°C. The reaction was quenched by the addition of sample buffer comprising 62.5 mM Tris– HCl, pH 6.8, 2% SDS, 5% β-mercaptoethanol, 0.08% bromophenol blue, and 30% glycerol and boiling of the sample before analysis of the products by 12 % SDS polyacrylamide gel electrophoresis. The gels were visualised either by Coomassie Blue staining or by immunoblotting using anti-ubiquitin antibodies and development in a Biorad ChemiDoc system. A mouse anti-DUB16 antibody (**Supplementary Figure 2**) was used for immunodetection of the recombinant enzyme.

### Cleavage of Ub-L40

10 μg of purified Ub-L40 substrate was incubated with 30 ng of DUB16 at 24°C and reaction aliquots were removed over the time range 0 - 32 minutes and quenched in SDS-PAGE sample buffer. The reaction products were resolved by denaturing 17.5 % polyacrylamide gel electrophoresis and visualised by staining with Coomassie blue dye.

### Protein crystallisation and structure determination

For structural studies, we recloned the DUB16 coding sequence so that enzyme with a smaller tag could be purified. For this purpose, we amplified by PCR the coding sequence of DUB16 from *L. donovani* LV9 genomic DNA template using the primers DUB16For and DUB16Rev (**Supplementary Table 2**). These incorporate restriction enzyme recognition sequences for cloning into the *Nde*I and *Hin*dIII sites of the plasmid pBSK2. This construct gives rise to an N-terminally HRV-3C protease cleavable His-tagged DUB16. DUB16 from the BPK and LV strains of *L. donovani* are identical. Digestion with 3C protease gives full length DUB16 with three amino acids (Gly-Pro-His) appended to its N-terminus.

Crystallisation experiments were performed in MRC-Wilden 96 well plates using commercial screens set up using Hydra 96 and Mosquito liquid handling systems to dispense well and drop solutions respectively. From initial screens, small crystals of DUB16 appeared in drops containing ammonium sulphate as a precipitant. Following a buffer pH screen and optimisation of protein and precipitant concentrations, crystals suitable for structure determination were grown in sitting drops composed of equal volumes of protein at 16 mg.ml^-1^ in 150 mM NaCl, 25 mM Tris-HCl pH 8.0, 1 mM DTT and a crystallisation solution of 1.9 M ammonium sulphate in 100 mM bis-Tris propane buffer (pH 7.0). For X-ray diffraction data collection, a single crystal was transferred to a solution of the mother liquor supplemented with 25% ethylene glycol as a cryoprotectant.

X-ray diffraction data were collected on beamline i03 at the DIAMOND Light Source (**Table 1**) and the data were processed using the xia2 suite. The point group is P321 with systematic absences suggesting a space group of either P3_1_21 or P3_2_21 with the cell dimensions indicating the presence of two molecules in the asymmetric unit giving a solvent content of 54%. The structure was solved by Molecular Replacement using the coordinates for human UCHL3 (PDB 1UCH, 37% sequence identity) as a search model in the programme MOLREP (47). Two solutions were found and the space group was unambiguously assigned as P3_1_21. Initial model refinement reduced the starting R factor from 52% to 42%. The model was rebuilt in the programme BUCCANEER (48) reducing the R factor to 23%. After cycles of manual building in COOT (49) and refinement using REFMAC (50), the *R*- and *R*-free factors for the final model are 21.1% and 23.4%. In the B molecule residues 162-166 are disordered while in the A molecule residues 127-130, 160-167 and 219-223 are disordered.

### Size Exclusion Chromatography with Multi-Angle Laser Light Scattering (SEC-MALLS)

A 100 μl DUB16 sample at 3.5 mg.ml^-1^ in 20 mM HEPES pH 7.5, 500 mM NaCl was resolved on a Superdex S75-210/300 GL column (GE Healthcare) run at 0.5 ml.min^-1^ with an HPLC system (Shimadzu). Light scattering data were collected continuously on material eluting from the column using an in-line Wyatt Dawn Heleos LS detector with an inline Wyatt Optilab rEX refractive index detector and an SPD-20A UV detector. Molecular weights were calculated by analysing data with the Wyatt program ASTRA.

### Thermal Shift Assay

Unfolding of protein DUB16 (12.5 μM) in the presence and absence of a 2- fold excess of compound IMP-1711 was measured by the change in fluorescence of SYPRO Orange dye (Sigma) with a Stratagene Mx3005P instrument. DMSO (final concentration of 5%) was used as a control. Data were analysed using JTSA software (http://paulsbond.co.uk/jtsa/).

### Generation of DUB16 overexpressing strains

*L. mexicana* (MNYC/BZ/62/M379) promastigote forms were grown at 25°C in Medium 199 (Gibco) supplemented with 10% (v/v) heat-inactivated fetal calf serum (hi-FCS) (Gibco) and 1% (v/v) penicillin/streptomycin solution (Sigma-Aldrich). Mid-log cultures were used for transfections with 10 μg of pNUS::DUB16::NEO (pGL 2831) (10) per 10^7^ promastigote cells. Transfection was performed by electroporation using an Amaxa Nucleofector 4D program FI-115 and the Unstimulated Human T-Cell Kit. Where required, parasites were grown with the selective antibiotic G418 (Neomycin) at 50 μg ml^-1^. Antibiotics were sourced from Invivogen.

### Activity based probe assay

Deubiquitinase activity-based probe assays were conducted as previously described (10), with the following modification. Instead of a Cy5-Ubi-PA probe, a HA-Ahx-Ahx-Ub-PA probe was used (UbiQ, Amsterdam). This contains a HA peptide linked to the N-terminus of Ubiquitin by two aminohexanoic acid groups, the ubiquitin possesses a C-terminal propargylamide warhead (UbiQ-078). During cell lysis the previous protease inhibitor mix was replaced with a defined commercial cocktail - HALT (Thermo Fisher). After HA-Ubi-PA labelling the reaction mix was stopped with the addition of 4x LDS NuPAGE buffer (Thermo Fisher) and separated using 4-20% Bio-Rad TGX Stain-Free gels with tris-glycine-SDS running buffer at 200 V for 75 minutes. Proteins were visualised in the gel by activating the Stain-Free system using a ChemiDoc MP for 45 seconds, followed by rapid auto-exposure. Proteins were then transferred to PVDF membranes with an iBlot 2 system (Thermo Fisher) using preset P0. Membranes were blocked with 5% milk powder in TBST buffer followed by antibody labelling with anti-HA.11 (Clone 16B12, BioLegend, 1:5000) and anti-mouse HRP (Promega, 1;5000). HRP activity was detected using Clarity ECL substrate (Bio-Rad). The chemiluminescent signal was quantified using Image Lab software Version 6.1.0 build 7 Standard Edition (Bio-Rad) and plotted in Prism Version 10.3.1 (464) (GraphPad). Activity based-probe assays were conducted in triplicate.

### Leishmania Promastigote Cell Viability Assay

The dose-response profile of IMP-1710 (37) was conducted over a 5-day incubation period with the addition of resazurin for the last 8 hours of the assay as previously described (38). Resorufin signal was measured using a BMG LABTECH CLARIOstar microplate reader, data were subsequently processed and plotted in GraphPad Prism 10 Version 10.3.1 (464). Dose-response curves were fitted using the mode: log(inhibitor) *versus* normalized response -- Variableslope. Three biological replicates were conducted, each containing 3 technical replicates.

### Mass Spectrometry

Ub-L40 samples, before and after, treatment with DUB16 were diluted in aqueous 50% acetonitrile containing 1% formic acid by ESI-MS using direct infusion at 3 ml/min and a Bruker maXis qTOF mass spectrometer. Acquired mass spectra were summed for each spectrum and deconvoluted to neutral average mases using MaxEnt with a resolution setting of 1K-5K.

## Acknowledgements

We thank Sam Hart, Johan Turkenburg and the Diamond Light Source for access to beamline i03 (proposal number mx-24948), Ed Tate for providing inhibitors, Zach Armstrong for crystal handling and Adam Dowle, Andrew Leech and Chris Taylor of the Bioscience Technology Facility for technical assistance. This work was funded by the United Kingdom Grand Challenges Research Funde under grant agreement ‘A Global Network for Neglected Tropical Diseases’ grant number MR/P027989/1 and the Wellcome Trust (200807/Z/16/Z & 223045/Z/21/Z). EN is supported by a BBSRC Studentship as part of the White Rose Doctoral Training Partnership (BB/M011151/1).

## Author Contributions

**Methodology / Investigation / Formal analysis / Validation** – JAB, MK, NGJ, EMN, EJD, SAE

**Writing – original draft** - JAB, MK, AJW

**Writing – reviewing and editting** All authors

**Funding Acquisition / Resources / Conceptualisation / Supervision / Project administration** – NA, JCM, AJW

**Visualisation** – JAB, MK, NGJ, EMN

**Data Curation / Software** – N/A

---

**Supplementary Table 1.** Pairwise sequence identities among UCH-type deubiquitinases.

|  | **LdDUB16** | **LdDUB15** | **HsUCHL1** | **HsUCHL3** | **HsUCHL5** | **HsBAP1** |
| --- | --- | --- | --- | --- | --- | --- |
| **LdDUB16** | - | **26** | **33** | **37** | **25** | **26** |
| **LdDUB15** |  | - | **24** | **25** | **41** | **34** |
| **HsUCHL1** |  |  | - | **55** | **23** | **24** |
| **HsUCHL3** |  |  |  | - | **29** | **24** |
| **HsUCHL5** |  |  |  |  | - | **47** |
| **HsBAP1** |  |  |  |  |  | - |

Sequence identities among the UCH type DUBs of *L. donovani* and the human host. The sequences used in the alignment were LdDUB16 (UNIPROT Accession A0A504WVB8) residues 1-233; LdDUB15 (A0A3S7WY30) residues 1-230; HsUCHL1 (P09936) residues 1-223; HsUCHL3 (P15374) residues 1-230; HsUCHL5 (Q9Y5K5) residues 1-228; HsBAP1 (Q92560) residues 1-240.

**Supplementary Table 2.**
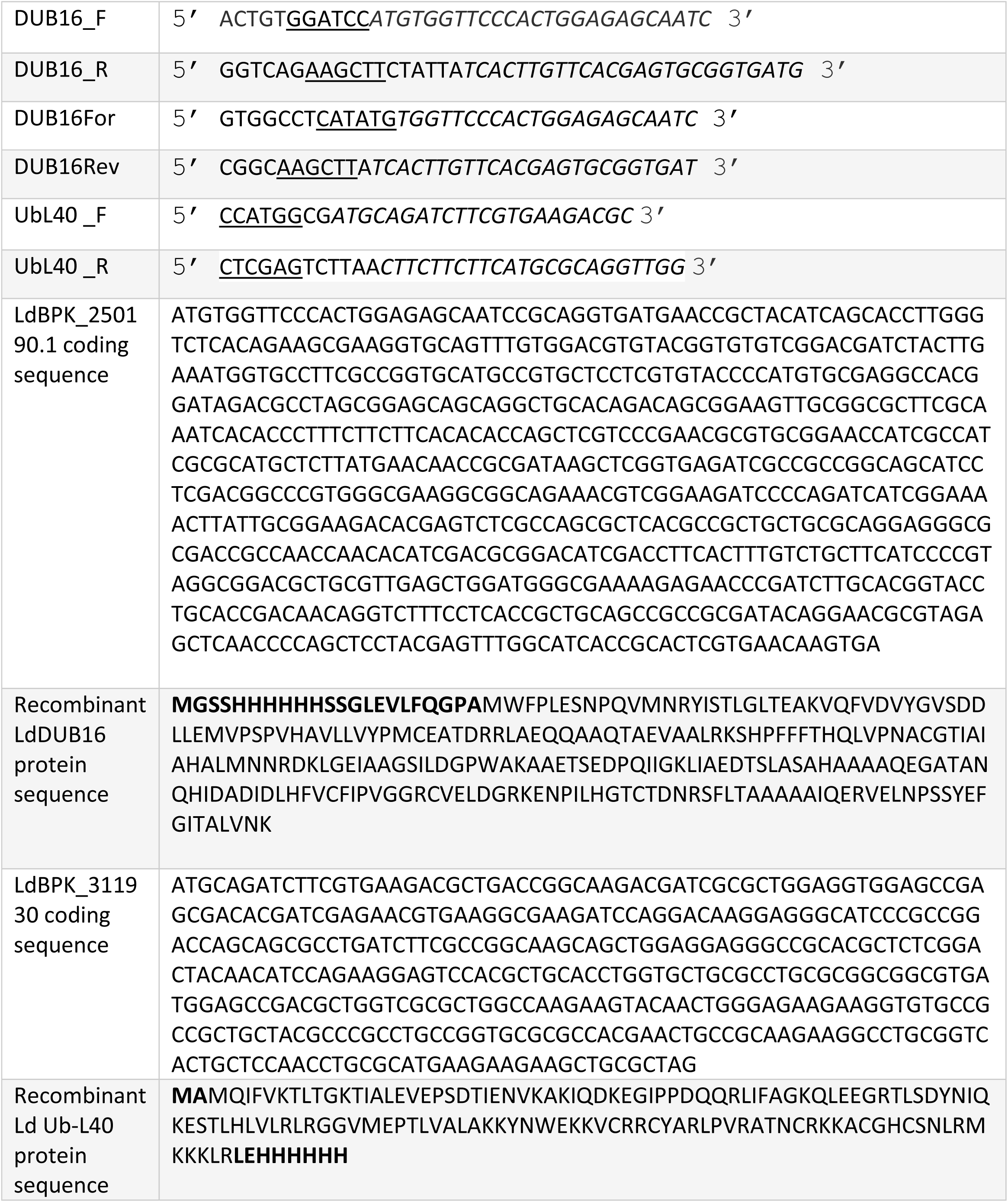
Oligonucleotide primers used in this work and amplified coding sequences.

**Supplementary Table 3.** Dose-Response Results.

|  | M379 | DUB16A | DUB16B |
| --- | --- | --- | --- |
| log(inhibitor)vs. normalized response-- Variable slope |  |  |  |
| Best-fit values |  |  |  |
| LogIC<sub>50</sub> | 0.3373 | 0.5624 | 0.5068 |
| HillSlope | -3.873 | -4.432 | -3.564 |
| IC<sub>50</sub> | 2.174 | 3.651 | 3.212 |
| 95% CI (profile likelihood) |  |  |  |
| LogIC<sub>50</sub> | 0.3083 to 0.3679 | 0.5378 to 0.5847 | 0.4687 to 0.5428 |
| Hill Slope | -5.192 to -3.097 | -9.972 to -3.383 | -4.871 to -2.791 |
| IC<sub>50</sub> | 2.034 to 2.333 | 3.450 to 3.844 | 2.942 to 3.490 |
| Goodness of Fit |  |  |  |
| Degrees of Freedom | 79 | 79 | 79 |
| R squared | 0.9516 | 0.9613 | 0.9321 |
| Sum of Squares | 7666 | 6142 | 10686 |
| Sy.x | 9.851 | 8.817 | 11.63 |
| Number of points |  |  |  |
| # of X values | 81 | 81 | 81 |
| # Y values analyzed | 81 | 81 | 81 |

**Supplementary Figure 1.**
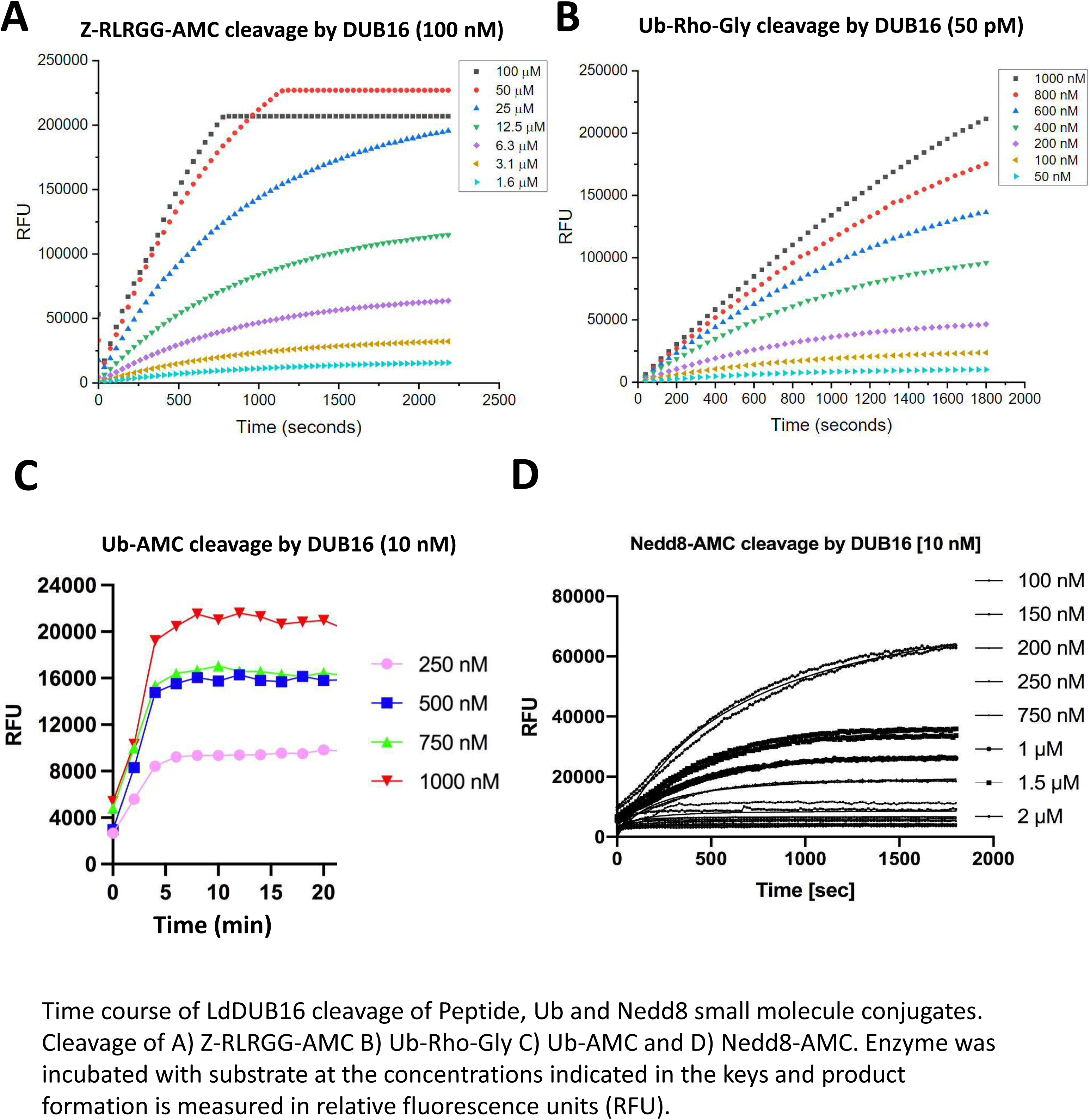
Time courses of the DUB16 catalysed cleavages of A) Z-RLRGG-AMC B) Ub-Rho-Gly C) Ub-AMC and D) Nedd8-AMC.

**Supplementary Figure 2.**
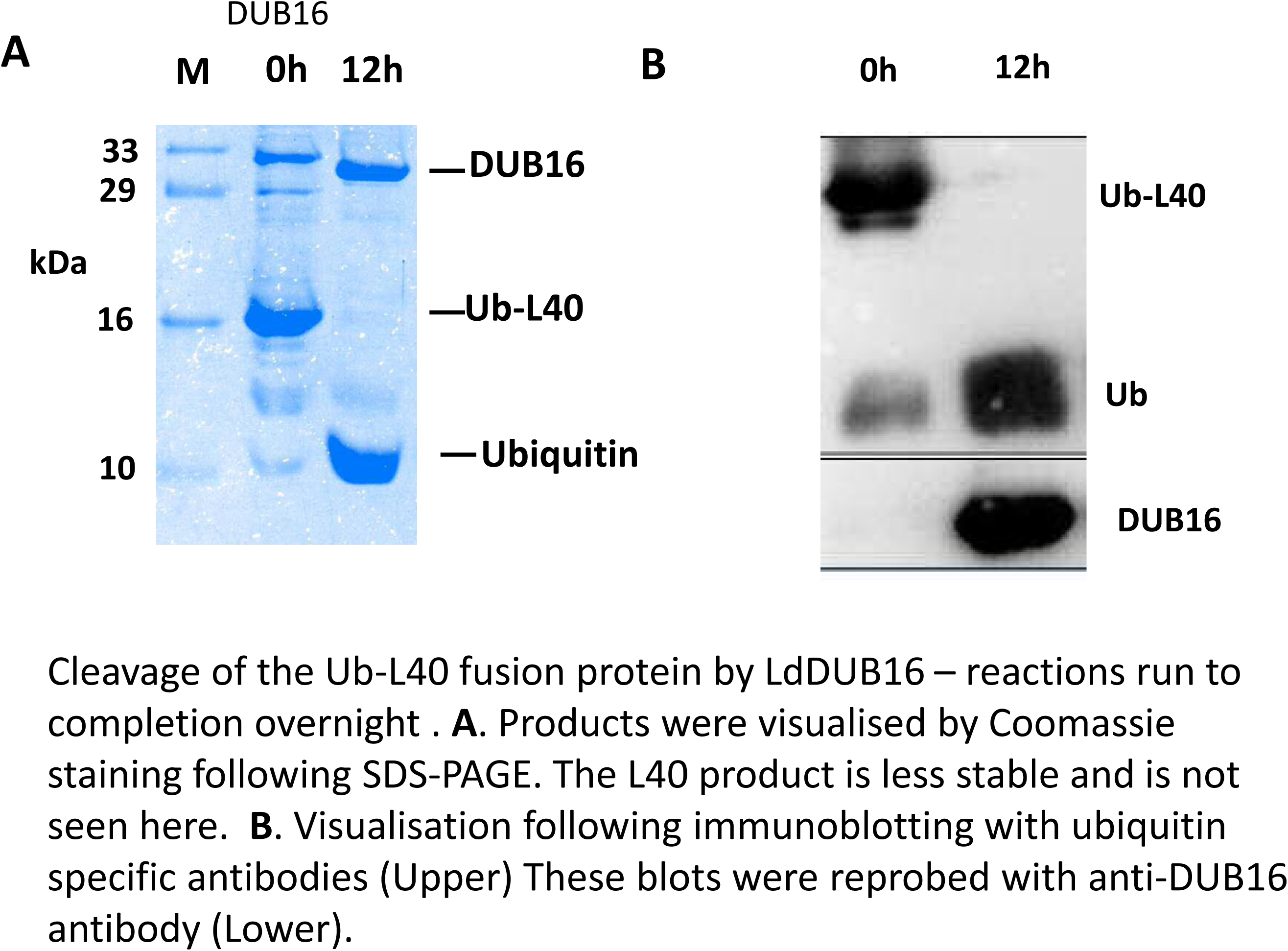
Cleavage of the Ub-L40 fusion protein by LdDUB16

**Supplementary Figure 3.**
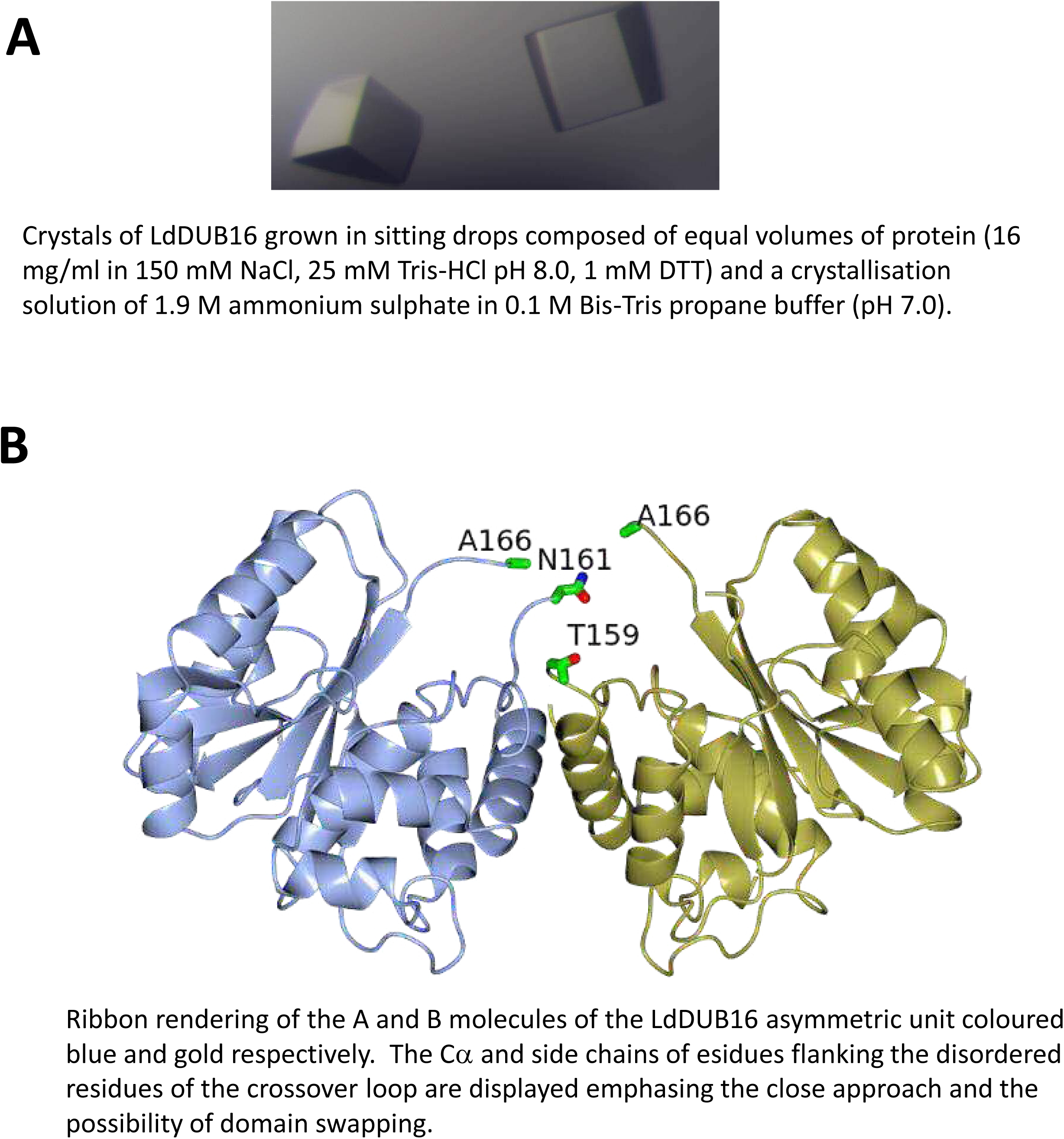
Crystals of DUB16 and the two chain of the asymmetric unit.

**Supplementary Figure 4.**
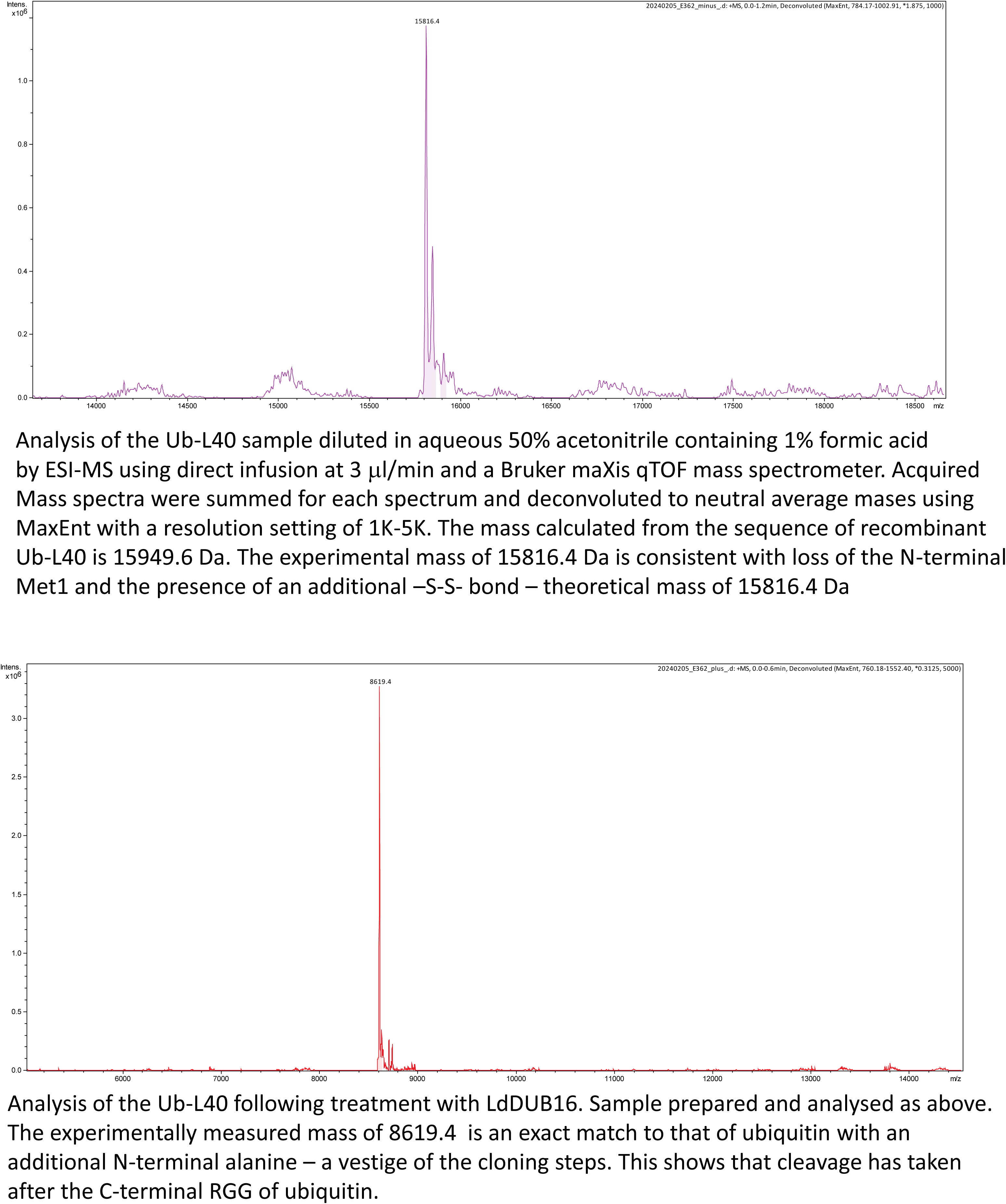
Mass spectrometry data

## Notes

### Competing Interest Statement

The authors have declared no competing interest.

